# Unveiling the complete invasion history of *D. melanogaster*: three horizontal transfers of transposable elements in the last 30 years

**DOI:** 10.1101/2024.04.25.591091

**Authors:** Riccardo Pianezza, Almorò Scarpa, Anna Haider, Sarah Signor, Robert Kofler

**Author notes:** Contributed equally (Riccardo Pianezza, Almorò Scarpa).

## Abstract

Transposable elements (TEs) are repetitive sequences capable of mobilizing within genomes, exerting significant influence on evolution throughout the tree of life. Using a novel approach that does not require prior knowledge about the sequence of repeats, we identified three novel TE invasions in *D. melanogaster* : *MLE* spread between 1990-2000, *Souslik* between 2009-2012, and *Transib1* between 2013-2016. We recapitulate prior findings, revealing that a total of 11 TEs invaded *D. melanogaster* over the past two centuries. Leveraging data from over 1400 arthropod genomes, we provide evidence that the TE invasions were triggered by horizontal transfers, with *D. simulans* and species of the *D. willistoni* group acting as putative donors. Through analysis of *∼*600 short-read datasets spanning diverse geographic regions, we reveal the rapidity of TE invasions: *Transib1* swiftly multiplied from three isolated epicenters in 2014 to all investigated populations within just two years. Our findings suggest that anthropogenic activities, facilitating habitat and population expansions of *D. melanogaster*, might have accelerated the rate of horizontal transposon transfer as well the spread of the TEs into the worldwide population. Given the significant impact of TEs in genomes and the potential involvement of humans in their dispersal, our research has crucial implications for both evolution and ecology.

## Introduction

Human activity is reshaping the global distribution of species at unprecedented rates [Turbelin et al., 2017], with climate change emerging as a significant consequence of anthropogenic influence, thereby reshaping species habitats [Gutierrez and Ponti, 2022, Ward and Masters, 2007, Ma and Ma, 2022]. As species move into new environments so do their pathogens, changing the assemblage of organisms that are now being exposed to these pathogens [Feurtey et al., 2023, Bebber et al., 2013, Anderson et al., 2004]. Transposable elements (TEs) are intra-genomic parasites that may also be affected by these dynamics. As species distributions change, TEs have the opportunity to invade genomes that were previously unavailable to them and shape the genomic evolution of a broad group of organisms.

TEs are short stretches of DNA that are able to copy themselves to new genomic locations, increasing their abundance in the genome [Hickey, 1982]. Some TEs have been shown to be beneficial to the host, such as insertions conferring insecticide resistance in *Drosophila* [Mateo et al., 2014, Aminetzach et al., 2005, Casacuberta and González, 2013]. However, many TEs have deleterious effects, for example due to the direct effects of the insertions or ectopic recombination among dispersed insertions [Nuzhdin, 1999]. To suppress their movement, hosts have developed dedicated defense mechanisms frequently involving small RNAs [Sarkies et al., 2015]. For example, in *Drosophila*, small RNAs, termed piRNAs, suppress TEs transcriptionally and post-transcriptionally [Brennecke et al., 2007, Gunawardane et al., 2007, Le Thomas et al., 2013, Sienski et al., 2012]. TEs that are silenced by the host will gradually accumulate mutations, ultimately leading to their immobilization and potential loss over time. However some TEs families, like non-LTR families, are predominately vertically transmitted [Malik et al., 1999], suggesting that they can escape this gradual decay, possibly by maintaining a low level of activity which preserves their sequence over time. Alternatively, other TE families may escape potential extinction by horizontal transfer to a naive species not having the TE [Peccoud et al., 2017].

Horizontal transfer of TEs amongst insects has been observed to be widespread [Peccoud et al., 2017, Pianezza et al., 2024, 2023, Kofler et al., 2015, Daniels et al., 1990]. Most evidence for horizontal transfer is based on indirect evidence, such as a high sequence similarity among insertions of different species [Peccoud et al., 2018, Wallau et al., 2012]. Direct observations of TE invasions have rarely been documented. The first well-documented case is the invasion of the *P* -element in *D. melanogaster* between 1950-1980, following horizontal transfer from *D. willistoni* [Kidwell, 1983, Anxolabéhère et al., 1988, Daniels et al., 1990]. This event was solely possible because *D. melanogaster* extended its habitat into the Americas about 200 years ago, thereby establishing contact with *D. willistoni*, a species that is endemic to South and Central America [Keller, 2007, Engels, 1983]. Later it was shown that the *I* -element, *Hobo* and *Tirant* also spread in *D. melanogaster* populations between 1930 and 1960 [Schwarz et al., 2021, Kidwell, 1983, Periquet et al., 1989, Daniels et al., 1990, Bucheton et al., 1992, Bonnivard et al., 2000]. Analysing the genomes of historical *D. melanogaster* specimens from museum collections, we showed that three additional TEs —*Blood, Opus*, and *412* — spread in *D. melanogaster* between 1850 and 1930 [Shpak et al., 2023, Scarpa et al., 2023]. Based on repeat libraries generated for different *Drosophila* species, we recently showed that *Spoink* invaded *D. melanogaster* between 1983 and 1993 [Pianezza et al., 2023], raising the number of TEs that spread in *D. melanogaster* to eight. Here we asked if additional TEs, that are not found in any of the extant repeat libraries, could have invaded *D. melanogaster* populations. In order to identify all recent TE invasions, we need to employ an approach that does not rely on any repeat library. We reasoned that TE invasions should lead to a recognisable pattern, such that sequences present in the genomes of recently collected strains ought to be absent in older ones. Using this approach, we indeed found that three additional TEs invaded natural *D. melanogaster* populations during the last 30 years.

We show that *MLE* (*Micropia*-like-element), a LTR retrotransposon of the *gypsy* /*mdg3* superfamily, spread in *D. melanogaster* between 1990 and 2000, *Souslik*, a LTR retrotranspsoson of the *gyspy* /*osvaldo* superfamily, between 2009 and 2012 and *Transib1*, a DNA transposon, between 2013 and 2016. We show that the *MLE* invasion was triggered by a horizontal transfer from a species of the *willistoni* group, whereas the *Transib1* and *Souslik* invasions were triggered by a horizontal transfer from the closely related *D. simulans*. Based on 585 samples collected during the invasions from different geographic regions, we were able to trace the spatio-temporal dispersal of *Transib1* . The invasion of *Transib1* commenced in at least three geographically isolated epicenters around 2014 (France, Ukraine, North America). Within a mere two years, *Transib1* spread to all investigated populations, demonstrating that TEs may rapidly infect geographically distinct populations.

## Results

### *MLE, Souslik* and *Transib1* recently spread in the *D. melanogaster* genome

Using multiple repeat libraries, we previously discovered that the TE *Spoink* invaded *D. melanogaster* between 1983 and 1993 [Pianezza et al., 2023]. This made us wonder whether other TEs, that are either absent or incompletely represented in the available repeat libraries, could have spread in *D. melanogaster* more recently. A major limitation in the identification of novel invasions is the need for a library of known TEs. We sought to circumvent this by developing an approach which can identify new TE insertions without a reference library. We reasoned that short reads from an old strain (before an invasion), aligned to an assembly of a recently collected strain (after an invasion), should result in coverage gaps at insertion sites of novel TEs (fig. 1A; supplementary figs S1, S2). To detect signatures of novel invasions, we developed a software called GenomeDelta which aligns short reads, identifies coverage gaps (corresponding to the individual insertions of a novel TE family), extracts their sequences and clusters them by similarity (Pianezza et al. in preparation). Here, we aligned reads of *Canton-S*, which was sampled in 1935, to the assembly of *Tom008*, a strain collected in 2016 [Lindsley and Grell, 1968, Rech et al., 2022]. We identified several sequences which were causing coverage gaps, many of which serve as a proof-of-concept as they corresponded to known invasions that occurred after 1935, such as for the *I* -element, *Tirant* and *Spoink* (supplementary fig. S1). Interestingly, we discovered three additional TE sequences leading to coverage gaps (fig. 1A; supplementary fig. S1).

The first sequence corresponds to a *Micropia*-like element (henceforth *MLE*) that was originally identified in two assemblies from the *Drosophila* Genetic Reference Panel, which were sampled in 2003 in Raleigh (GenBank MN418888; [Ellison and Cao, 2020, Mackay et al., 2012]). As insertions of *MLE* were absent in the reference strain *Iso-1*, the authors suggested that *MLE* recently invaded the *Drosophila* Genetic Reference population [Ellison and Cao, 2020]. *MLE* is a 5360 bp LTR retrotransposon with an LTR of length 361 bp. It has roughly 70% similarity with the consensus sequence of *Micropia* annotated in *D. melanogaster* [Quesneville et al., 2005]. Based on the alignment of the reverse transcriptase domain, *MLE* is a member of the *gypsy* /*mdg3* group, similarly to *Spoink* (supplementary fig. S3; [Kapitonov and Jurka, 2003, Pianezza et al., 2023]). *MLE* and *Spoink* have a different length (5360 vs. 5216 bp), different sized LTRs (361 vs. 349 bp) and generally share little sequence similarity (63% identical). In addition, *MLE* and *Spoink* insertions are at different genomic sites (supplementary fig. S4).

**Figure 1:**
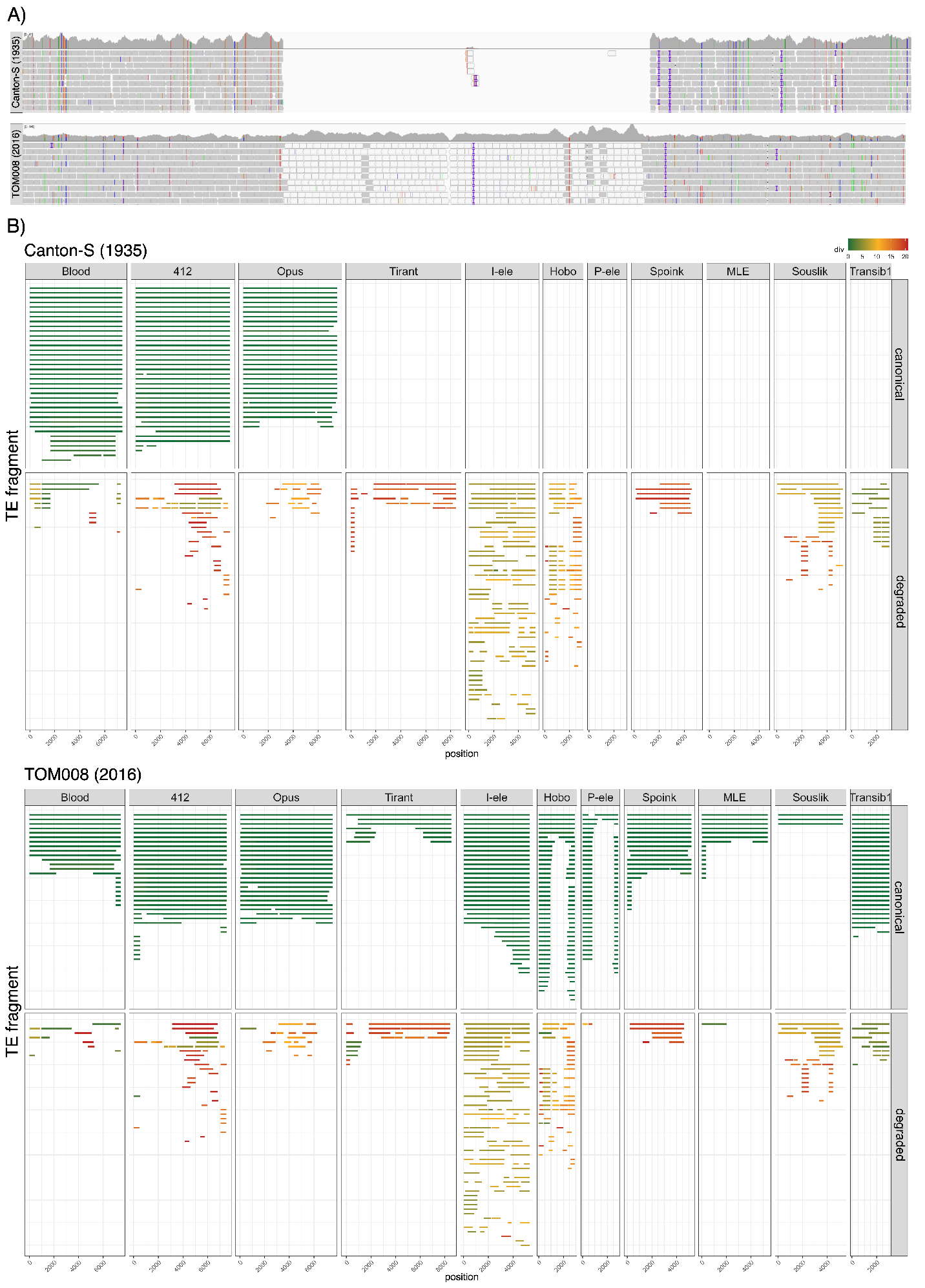
Overview of the horizontal transfers of TEs in *D. melanogaster* during the last two centuries from long-read assemblies. A) Illustration of our approach to find TE invasions. A novel TE invasion can be detected by gaps in the coverage, when reads of an old strain (e.g. *Canton-S*, sampled 1935) are aligned to the assembly of a recent strain (e.g. Tom008; sampled 2016; top panel). The gap in the example is due to *Souslik* . By contrast, no coverage gap is found if reads from the same strain are aligned (bottom panel). B) Horizontal bars represent the matching regions relative to the consensus sequence of the TE. Colors represent divergence from the consensus sequence. Based on a divergence threshold of 1.5%, we classified the fragments into canonical (*≤*1.5%) and degraded (*>*1.5%). Note that canonical insertions are present in the recent but absent in the old strains. The first three TEs spread before 1935, which explains the presence of canonical insertions in both assemblies.

The sequence of the second coverage gap shares 99.68% sequence similarity with *Souslik*, a TE previously described in *D. simulans* [Glukhov et al., 2016]. *Souslik* has a length of 5275 bp and an LTR of 210 bp. *Souslik* is a member of the *Ty3* /textitGypsy group and enocdes a *gag* -*pol* fusion product, but no *env* [Glukhov et al., 2016]. The phylogenetic tree based on the reverse transcriptase domain suggests that *Souslik* is member of the *gyspy* /*osvaldo* group (supplementary fig. S3). However, previous work suggested that *Souslik* may be more closely related to *Mag* elements [Glukhov et al., 2016] (which are not represented in the phylogenetic framework of *D. melanogaster* LTR retrotransposons [Kapitonov and Jurka, 2003]).

Finally, the third sequence has a high similarity with *Transib1* in *D. melanogaster* (96.06% over 72% of the length) and *D. simulans* (99.93% over 100 % of the length) [Kapitonov and Jurka, 2003]. *Transib1* is a DNA transposon of length 3030 bp. Note that the sequence of *Transib1* in *D. melanogaster* was reconstructed from fragments of the transposon that were dispersed throughout the genome [Kapitonov and Jurka, 2003]. Prior to this work, no full-length insertion of *Transib1* in *D. melanogaster* was described [Kapitonov and Jurka, 2003].

To further investigate these recent invasions, we analysed TE insertions in long-read assemblies of *D. melanogaster* strains collected at different times during the last century (*Canton-S* 1935 and *Tom008* 2016). Using RepeatMasker, we identified sequences with similarity to any of the 11 TEs that may have invaded *D. melanogaster* recently (fig. 1B). We refer to insertions with a high similarity to the consensus sequence (*≤*1.5%) as canonical insertions, to differentiate them from degraded copies. As expected, in *TOM008* (2016) we found canonical insertions for all eight TEs that were previously shown to have spread in *D. melanogaster* in the last two centuries. Canonical insertions were absent in *Canton-S* (1935) for the five TEs that spread after 1935, i.e. *Tirant, I* -element, *Hobo, P* -element, and *Spoink* . Since *Blood, Opus* and *412* spread between 1850 and 1933, canonical insertions of these TEs were present in both strains (*Canton-S* and *Tom008* [Scarpa et al., 2023]. No high quality long-read based assembly is available for any strain sampled before 1925-1935 (i.e. before *Canton-S* and *Oregon-R* were sampled). Importantly, we found canonical insertions of *MLE, Souslik* and *Transib1* in *TOM008* (2016) but not in *Canton-S* (1935; fig. 1B). Degraded fragments were found for all TEs, except for the *P* -element and *MLE*, in both *TOM008* and *Canton-S* (fig. 1B). Ancient invasions of TEs sharing some sequence similarity with the focal TE are the most probable source of these degraded fragments. An investigation of the insertion sites reveals that the degraded fragments of the TEs are close to centromeric heterochromatin, whereas the canonical insertions are distributed over the entire chromosome arm (supplementary fig. S4). An analysis of the assemblies of multiple strains collected between 1968 and 2015 confirms that the canonical insertions of *MLE, Souslik* and *Transib1* gradually emerged during the last decades, while degraded fragments are present in all analysed strains (supplementary fig S5). Next, we analysed the copy number of all TEs in several strains sampled around 1980 and 2015 with DeviaTE [Weilguny and Kofler, 2019]. For each TE family, DeviaTE normalizes the average coverage of a TE to the average coverage of single copy genes, which allows inferring the TE copy number per haploid genome [Weilguny and Kofler, 2019]. An analysis of the read coverage shows that *MLE, Souslik* and *Transib1* have a uniformly elevated coverage in strains sampled in 2016 as compared to strains sampled before 1968 (supplementary figs. S6, S7, S8). For *Souslik* and *Transib1*, a low level of coverage based on diverged reads can also be found in strains sampled before 1968, in agreement with degraded fragments being present in all analysed strains (supplementary figs. S6, S7, S8).

Finally, investigating the population frequency of the TE insertions, we found that canonical insertions are largely segregating at low frequency, whereas degraded insertions are mostly fixed (supplementary fig. S9). This is consistent with the hypothesis that canonical insertions are recent whereas degraded insertions are more ancient.

In summary, we suggest that the LTR-retrotransposons *MLE* and *Souslik*, as well as the DNA transposon *Transib1*, recently spread in the *D. melanogaster* genome. Degraded fragments, likely remnants of past invasions, can be found in all analysed strains for *Souslik* and *Transib1*, but not for *MLE* .

### History of TE invasions in *D. melanogaster* during the past two centuries

Given that our unbiased approach (GenomeDelta) likely detected all recent TE invasions, we can now reconstruct the complete history of TE invasions in *D. melanogaster* during the last two centuries. We analysed 585 short read datasets from individual strains or pooled populations collected during the last two centuries (supplementary file S1). We inferred the invasion status from two key metrics, i.e. the TE copy numbers and the frequency of SNPs diagnostic for canonical insertions. The copy numbers of TEs were estimated with DeviaTE [Weilguny and Kofler, 2019]. For each of the 11 TEs that recently spread in *D. melanogaster*, we identified a set of diagnostic SNPs, occurring in canonical but not in degraded insertions (supplementary fig. S10). Since we found that diagnostic SNPs of a given TE yield consistent frequency estimates, we averaged the frequency of the diagnostic SNPs for each TE (supplementary fig. S11). For TEs where we did not find diagnostic SNPs (i.e. no short reads aligning to the TE; for the *P* -element and *Opus*), we used frequency estimates of 1.0 (presence of reads) and 0.0 (absence of reads). The frequency of diagnostic SNPs roughly reflects the abundance of canonical insertions, where, for example, a frequency of 0.8 suggests that about 80% of the insertions of a given family are canonical and 20% degraded.

Consistent with previous works, both of our key metrics (copy number and diagnostic SNPs) suggest that *Blood, Opus* and *412* spread between 1850 and 1933, followed by *Tirant* (1933-1950), the *I* -element (1933-1950), *Hobo* (around 1950), the *P* -element (around 1960) and *Spoink* which spread between 1983-1993 (fig. 2; [Scarpa et al., 2023, Schwarz et al., 2021, Bucheton et al., 1992, Periquet et al., 1989, Bonnivard et al., 2000, Kidwell, 1983, Anxolabéhère et al., 1988, Daniels et al., 1990, Pianezza et al., 2023]). Our data further suggest that *MLE* spread between 1990 and 2000, slightly after *Spoink* (1983-1993; [Pianezza et al., 2023]). *Souslik* spread between 2009 and 2012 and *Transib1* between 2013 and 2016 (fig. 2).

**Figure 2:**
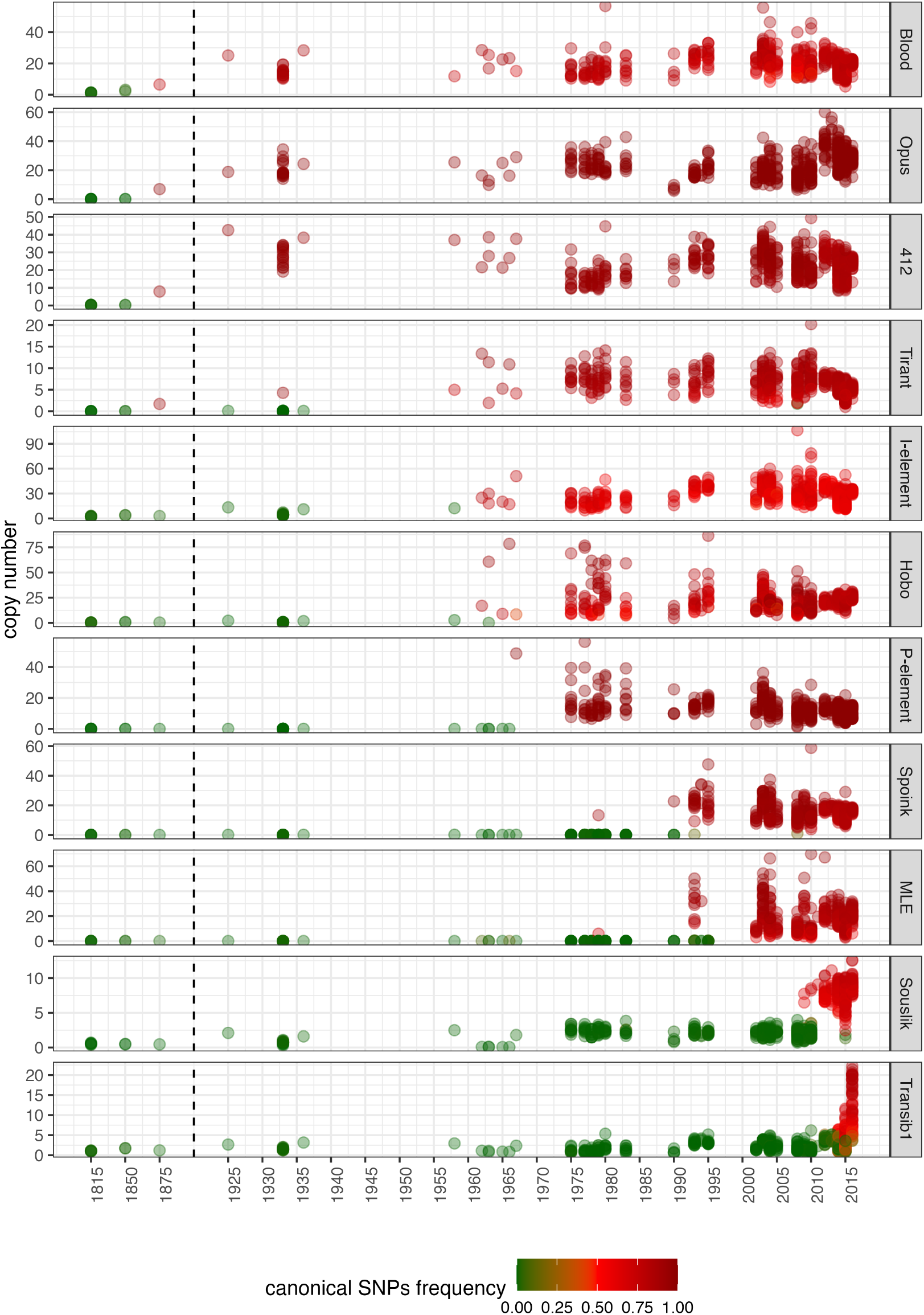
Complete invasion history of *D. melanogaster* during the last two centuries. We monitored the copy numbers of the TEs (y-axis) as well as the frequency of SNPs diagnostic for canonical insertions (color) in *D. melanogaster* strains and pooled populations sampled during the last two centuries. A recent invasion can be recognized by a sudden increase in TE copy numbers and the frequency of diagnostic SNPs.

Next, we traced the invasion of these 11 TEs in 49 long-read assemblies of *D. melanogaster* strains collected between 1925 and 2018 (supplementary file S2). A major advantage of long-read assemblies is that the copy number of canonical insertions can be directly inferred (from the RepeatMasker output). However, a drawback is the lower temporal resolution compared to short-read data, largely due to the limited number of high-quality assemblies. The abundance of canonical insertions in long-read assemblies reproduces the timing of the TEs previously shown to have spread in *D. melanogaster* between 1925 and 2018, i.e.: *Tirant, I* -element, *Hobo, P* -element and *Spoink* (supplementary fig S12). Interestingly, canonical *Tirant* insertions are still absent in a few recently collected strains (supplementary fig S12). No canonical insertions of *MLE, Souslik* and *Transib1* were found in any strain collected before 2002 (supplementary fig. S12). The first canonical *MLE* insertions are found in strains collected in 2003, for *Souslik* in 2011 and for *Transib1* in 2015. The abundance of canonical insertions in long-read assemblies is thus consistent with the timing of the invasion inferred from the short read data.

In summary, we inferred the complete invasion history for all 11 TEs that spread in *D. melanogaster* between 1810 and 2018. We were able to reproduce the timing of the eight TEs previously shown to have spread in *D. melanogaster* recently, and additionally showed that *MLE* spread between 1990 and 2000, *Souslik* between 2009 and 2012 and *Transib1* between 2013 and 2016

### Geographic spread of the invasions

Next, we examined the timing of geographic spread of the TEs across worldwide populations. Given the recent spread of *MLE, Souslik*, and *Transib1* over the last few decades, a wealth of resources is available to infer the timing of these invasions. In particular, consortia such as DrosEU (mostly Europe), DrosRTEC (mostly North America) and DGN (worldwide) contributed a vast number of short-read datasets of strains or pooled populations sampled from different locations during the last decades [Kapun et al., 2020, Machado et al., 2021, Kapun et al., 2021, Lack et al., 2015, 2016]. Using these rich resources, we investigated the spatiotemporal spread of *MLE, Souslik* and *Transib1*, relying on the diagnostic SNPs (fig. 3; supplementary fig. S13).

**Figure 3:**
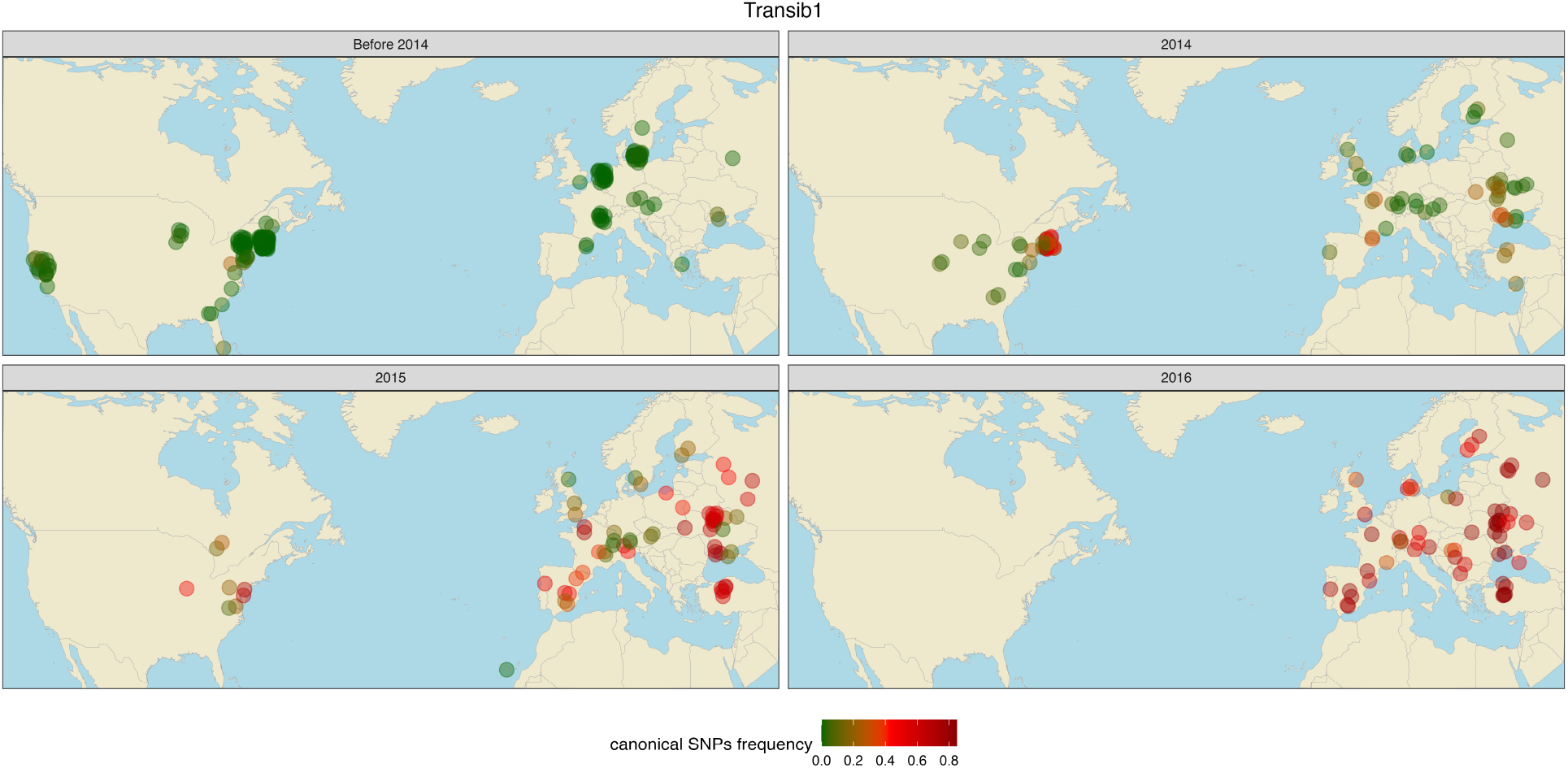
Geographic spread of canonical *Transib1* insertions in *D. melanogaster* samples (strains or pooled populations) collected from Europe and North America between 2013 and 2016. Colors indicate the frequency of SNPs diagnostic for the presence of the canonical *Transib1* insertions.

The first population in which we detected *MLE* was an African population collected in 1993, when it was not yet present in a population from China. After 1996, *MLE* was found in populations across the globe (supplementary fig. S13). Note that the early presence of *MLE* in Africa does not imply that the invasion started in Africa, as too few data from other populations are available between 1993-1995 to reliably infer the origin of the invasion from the spatio-temporal data (other lines of evidence suggest that the invasion started in South America; see below). Based on the abundance of canonical insertions in long-read assemblies, we confirm that *MLE* was absent in worldwide *D. melanogaster* strains before 1995 but present afterwards (supplementary fig. S14).

Global populations lacked *Souslik* insertions before 2009. *Souslik* was first found in North America in 2009 and had become widespread across most populations by 2011. However, in 2015 *Souslik* was still absent in populations sampled from the Caribbean archipelago of Guadeloupe (GS Sai 15 2 and GS Des 15 1; supplementary fig S13). This shows that geographic isolation of island populations may protect them, at least for some time, from TE invasions. The long-read assemblies confirm that canonical *Souslik* insertions were absent in global *D. melanogaster* strains sampled before 2003 and present afterwards (supplementary fig. S14).

The timing of the *Transib1* invasion was particularly fortunate, as it fell right into the time when DrosEU and DrosRTEC were collecting massive amounts of samples from Europe and North America (2013-2016; fig. 3). This allowed us to trace the *Transib1* invasion with unprecedented spatio-temporal resolution (fig 3). *Transib1* was largely absent in populations both from North America and Europe before 2014, with the exception of one sample from Virginia (VA ch 12 fall) collected in 2012. In 2014, a few *Transib1* copies were found in populations from Virginia, France and Ukraine. However, the vast majority of samples did not have *Transib1* insertions by 2014. In 2015, the *Transib1* invasion progressed rapidly, with *Transib1* insertions being found in many populations from both North America and Europe. By 2016, all sampled populations had *Transib1* insertions (fig 3). The spread of *Transib1* was extremely rapid. Within just two years (2014-2016), it spread from a few populations in Europe and North America to all the sampled populations. It is notable that the invasion did not spread from a single location, but started at multiple geographically isolated ‘epicenters’ (i.e. Ukraine, France, Virginia). Human activity (e.g. air traffic) may be responsible for this rapid spread of *Transib1* among isolated populations. The long-read assemblies confirm that canonical *Transib1* insertions are absent in all strains sampled before 2011 and present in strains sampled later (supplementary fig. S14).

In summary, we were able to trace the spatio-temporal spread of *MLE, Souslik*, and *Transib1* . The invasion of *Transib1* is particularly important, as it demonstrated that the spread of this TE occurred concurrently from several geographic locations and that within just two years the majority of the populations were infected. This extremely rapid dispersal of a TE has not been previously documented, as sampling of flies from the wild is generally not dense enough.

### Origin of *MLE, Souslik* and *Transib1*

The invasions of *MLE, Souslik* and *Transib1* in *D. melanogaster* were likely triggered by an horizontal transfer event from a different species. To identify the origin of the horizontal transfer, we investigated TE insertions in the assemblies of 266 drosophilids [Kim et al., 2021, 2023, Pianezza et al., 2024] (supplementary file S3), representing 242 species. Additionally, we analysed the reference genomes of 1226 arthropods species (supplementary file S4). We used RepeatMasker to identify *MLE, Souslik* and *Transib1* insertions in these genomes, together with the other 8 previously reported invaders. In agreement with previous works, we found insertions with a high similarity to *Blood, Opus, 412, Tirant*, the *I* -element and *Hobo* in *D. simulans* and related species, whereas the *P* -element and *Spoink* have similar insertions in species of the *willistoni* group [Pianezza et al., 2023, Daniels et al., 1990, Scarpa et al., 2023]. For *MLE*, we found highly similar insertions in species of the *willistoni, cardini* and *repleta groups*. Interestingly, *MLE* was entirely absent in *D. simulans* and other species of the *simulans* species complex (*D. mauritiana* and *D. sechellia*). For *Souslik* and *Transib1*, the most similar insertions were found in species of the *simulans* species complex. We did not find sequences with a high similarity to the 11 TEs that invaded *D. melanogaster* recently in the 1226 arthropods, outside of the drosophilids (supplementary fig. S15; apart from *MLE*, where insertions with some sequence similarity can be found in *Anastrepha obliqua* and *Merodon equestris*).

To further elucidate the source of the horizontal transfer, we generated phylogenetic trees of full-length insertions of *MLE, Souslik* and *Transib1* (fig. 4B). Insertions of all three TEs have short branches in *D. melanogaster*, consistent with a recent invasion (fig. 4B). The *MLE* insertions in *D. melanogaster* are nested in insertions of species of the *willistoni* group. This suggests that, much as the *P* -element and *Spoink*, a species of *willistoni* group acted as donor of *MLE* [Pianezza et al., 2023, Daniels et al., 1990]. Why species of the *willistoni* group repeatedly acted as donors to *D. melanogaster* is an open question. The phylogenetic trees of *Souslik* and *Transib1* suggest a different origin, as their insertions exhibit the closest relationship to *D. simulans*.

**Figure 4:**
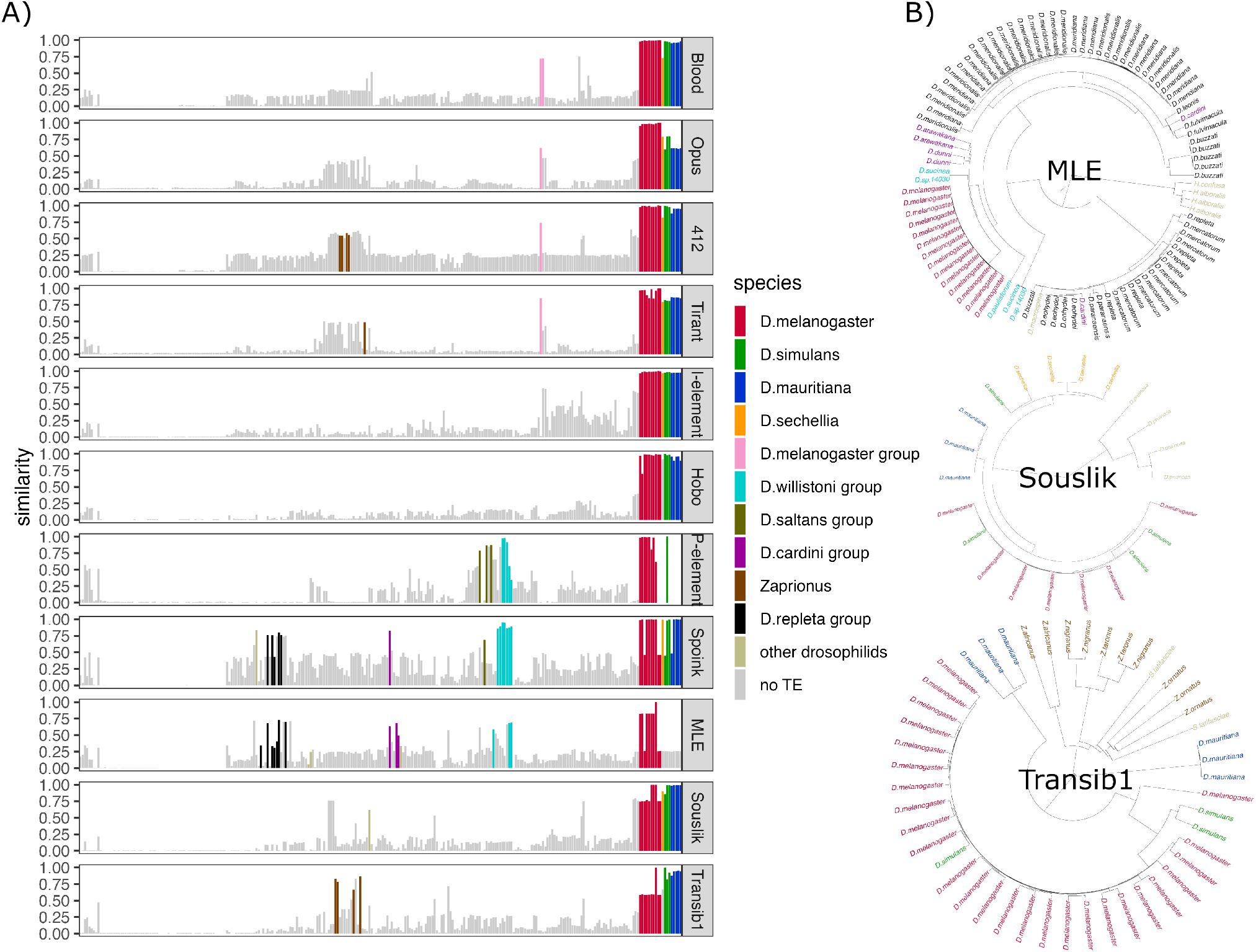
The *Souslik* and *Transib1* invasions were triggered by horizontal transfer from *D. simulans*, whereas *MLE* was transferred from a species of the *willistoni* group. A) Similarity of the 11 TEs that recently invaded *D. melanogaster* with insertions in 266 high-quality assemblies of 242 *Drosophila* species. The barplots show, for each species, the similarity between the given TE and the best match in an assembly. The species were arranged by relatedness. Colors indicate species or species groups with full-length insertions (*>*80% of length). B) Bayesian tree of *MLE, Souslik* and *Transib1* insertions in species having at least one full-length insertion.

In summary, our findings suggest that *MLE* invaded the genome of *D. melanogaster* through an horizontal transfer from a species of the *willistoni* group, while *D. simulans* was the donor of *Souslik* and *Transib1* .

## Discussion

Using a novel approach to identify recent genomic invaders independently of any repeat library, we found that three TEs, mle, *Souslik* and *Transib1*, spread in *D. melanogaster* populations between 1990 and 2016. All three invasions are more recent than the spread of *Spoink* (1983-1993) previously thought to be the most recent invasion [Pianezza et al., 2023]. The three invasions were probably triggered by a horizontal transfer from different species. Most previously reported cases of horizontal transfer of TEs were inferred from indirect evidence, such as an unusual high sequence similarity of the TE among species, a patchy distribution of the TE among closely related species and phylogenetic discrepancy between the TE and the host species [Peccoud et al., 2018, Wallau et al., 2012]. The three horizontal transfers described here are supported by much more direct and thus more compelling evidence, i.e. the absence of canonical insertions in old samples and their presence in recently collected samples (e.g. strains or pooled populations). This presence/absence pattern of the canonical insertions is supported by numerous short-read data and long-read assemblies (fig 2; supplementary fig S12).

Combined with the eight previously known cases, our work suggests that in total 11 TEs invaded *D. melanogaster* populations in the last two centuries following horizontal transfer [Scarpa et al., 2023, Schwarz et al., 2021, Bucheton et al., 1992, Periquet et al., 1989, Bonnivard et al., 2000, Kidwell, 1983, Anxolabéhère et al., 1988, Daniels et al., 1990, Pianezza et al., 2023]. Since our method for identifying novel invaders is unbiased, i.e. it does not require prior knowledge about the sequence of repeats, we argue that we identified all TE invasions in *D. melanogaster* populations between *∼*1810 (the earliest collected strain) and 2018 (the most recently collected long-read assembly). We may solely have missed invasions of TEs with a very high sequence similarity to resident TEs (which might not result in noticeable coverage gaps). It is not clear if horizontal transfer of a TE with high similarity to resident TEs could actually trigger an invasion, as the piRNA based host defence will likely be active against the resident TE and the piRNAs complementary to the resident TE will also target the novel TE (some mismatches are tolerated [Schwarz et al., 2021, Gainetdinov et al., 2023]). We may have further missed local invasions, that just happened in a few isolated subpopulations.

Although we inferred the complete invasions history between *∼*1810 and 2018, we think it is likely that additional TEs will invade *D. melanogaster* populations in the next decades. It is interesting to speculate which TEs may spread next. Given the high rate of horizontal transfer events between *D. melanogaster* and *D. simulans*, TEs absent in one of the two species could invade the other. For example, *Shellder* has spread in *D. simulans* and *D. mauritiana*, but it’s still absent in *D. melanogaster* [Ding et al., 2016] (Scarpa et al; in preparation). On the other hand we found *MLE* to be present in *D. melanogaster* but not in *D. simulans*. It is thus a promising candidate for next invasions in *D. simulans* and related species. It is also plausible that additional transposons will jump in *D. melanogaster* from species of the *willistoni* group.

In fact, one of the interesting patterns that has emerged is that the recent invasions in *D. melanogaster* are triggered by horizontal transfer either from *D. simulans* (*Blood, Opus, 412, Tirant, I* -element, *Hobo, Souslik, Transib1*) or a species of the *willistoni* group (*P* -element, *Spoink, MLE*). Horizontal transfer of TEs between the closely related *D. simulans* and *D. melanogaster* is not unexpected, as there could be a rare hybridization amongst some members of the complex. Additionally, the species have a somewhat overlapping ecology [Capy and Gibert, 2004] and could share potential vectors (e.g. viruses, parasites and *Wolbachia*) that might mediate the horizontal transfer of TEs. In agreement with this, previous works found horizontal transfer is most likely among closely related insect species [Peccoud et al., 2017]. However, *D. melanogaster* and species of the *willistoni* group are distantly related (100My [Obbard et al., 2012]. Thus, an important open question is why species of *willistoni* group repeatedly acted as donors of TE invasions in *D. melanogaster*, and not any of the many other species that came into contact with *D. melanogaster* following its habitat expansion (e.g. *D. pseudoobscura*, which is endemic in North America [Dobzhansky and Epling, 1944]). A related question is whether the horizontal exchange between *D. melanogaster* and species of the *willistoni* group is unidirectional or if species of the *willistoni* group also recently received TEs from *D. melanogaster* . It was for example suggested that Copia was recently horizontally transferred from *D. melanogaster* to *D. willistoni*, although recent works did not confirm this [Jordan et al., 1999, Rubin et al., 2011].

Given the amount of new horizontal transposon transfers in *D. melanogaster* in the last two centuries, another open question concerns the effect of these novel invasions on the genome and the phenotype. The 11 invasions increased the genome size of *D. melanogaster* by 0.8 - 0.91% during the last two centuries (supplementary table S1). They could also lead to inversions, translocations or other large genomic rearrangements, as they have in cichlids [Quah et al., 2024]. The 11 new TEs might thus have a substantial impact on the evolution of the genome of *D. melanogaster* . However, the invasions could also have notable phenotypic effects. Given that most TE insertions are likely deleterious to hosts [Elena et al., 1998, Pasyukova et al., 2004] it is plausible that the 11 TEs invasions had a negative impact on the fitness of *D. melanogaster* . Nevertheless, some insertions may also be beneficial [Casacuberta and González, 2013, González et al., 2008]. As the vast majority of the insertions of the 11 TEs are segregating at low frequency (supplementary fig. S9), the new TEs generated substantial standing genetic variation, which could act as substrate mediating rapid adaptation. For example, an insertions of the *P* -element, which spread between 1960-1980, contributed to insecticide resistance in *D. melanogaster* [Schmidt et al., 2010]. Overall, it is thus not clear whether these 11 invasion are a blessing or a curse. Future work investigating the distribution of fitness effects of novel insertions of these TEs may help to answer this question.

The high rate of recent invasions in the last two centuries is unlikely to be reflective of the overall background rate of TE invasion in *D. melanogaster* . *D. melanogaster* has about 130 TE families, of which 11 invaded during the last two centuries [Quesneville et al., 2005]. Assuming that we can identify TE families that are at least 2 million years old [Bergman and Bensasson, 2007], then one invasion every 20 years should result in about 100,000 families recognizable in the genome of *D. melanogaster* . Considering that each family invades in multiple waves (e.g repeated re-invasions as for example seen for the *I* -element, *Tirant, Blood*, etc), then we still ought to find *>*10,000 TE families. This discrepancy between the expected (*>*10,000) and observed (130) number of TE families suggest that the rate of invasion during the last two centuries is unusually high.

What is the source of such an unprecedented rate of horizontal transfer events during the last centuries? We think that the most plausible explanation is the recent habitat expansion of *D. melanogaster*, possibly mediated by anthropogenic activity. About 200 years ago, *D. melanogaster* spread from Europe and Africa to the Americas and Australia [A.H.Sturtevant, 1921, Keller, 2007, Bock and Parsons, 1981]. This habitat expansion brought *D. melanogaster* into contact with novel species that could act as donors of novel TEs. For example, the spread of *D. melanogaster* into the habitat of species of the *willistoni* group enabled the horizontal transfer of the *P* -element, *Spoink* and *MLE* . Habitat expansion will also result in an increase in population size and bring species into contact with novel vectors that could mediate the horizontal transfer of TEs, thus increasing the chances for horizontal transfer among species. Hence, the recent habitat expansion might also account for high rate of horizontal transfer between *D. melanogaster* and *D. simulans*, even though both species previously shared a common habitat for a long time (i.e. *>*200 years ago)

Another effect of anthropogenic activity could have been revealed by the spatio-temporal spread of *Transib1* . Strikingly, *Transib1* invaded almost all populations from Europe and North America in a mere two years (fig. 3). If we assume that *D. melanogaster* has about 15 generations per year [Pool, 2015], then *Transib1* penetrated *D. melanogaster* populations in 30 generations. Studies that monitored TE invasions in experimental populations of different *Drosophila* species revealed that it takes about 20 generations for TEs to spread in a population (i.e. to amplify from a few copies per individual to stable high copy numbers [Kofler et al., 2018, 2022, Selvaraju et al., 2024]). Although the spread in geographically distributed natural populations is likely much slower than in the panmictic experimental populations, it is plausible that a TE may reach high copy number in natural populations within 30 generations. It is likely that human activity facilitated the spread of *Transib1* across the different populations, for example by carrying flies as stowaways with cargo. The fact that the *Transib1* invasion commenced at three different geographically distant epicenters in 2014 is in favor of human activity (fig. 3). TEs would likely spread more slowly in the absence or with reduced human activity, as suggested by the *P* -element invasion. This invasion occurred during a period when air traffic, for example, was less intense than now (1960-1980) and took several decades to reach all sampled populations [Kidwell, 1983, Anxolabéhère et al., 1988].

Given that many other insects species are currently expanding their habitats due to human activity and climate change [Parvizi et al., 2023, Tang et al., 2019, Lee et al., 2011], it is a crucial question whether those other species also exhibit such a high rate of recent horizontal transposon transfers.

On this regard, two considerations are important. First, horizontal transfer is a stochastic event, and will likely not occur immediately upon contact between species. As a result, TE invasions will likely lag behind habitat expansions. Second, habitat expansion of insects lags behind the habitat expansion of plants (many insects depend on some host plants). Given the amount of habitat expansions of plants, it has been estimated that many more habitat expansions of insects are yet to follow [Bonnamour et al., 2023]. As a net effect of these two lag-events, the rate of TE invasions may be increasing for many insect species in the coming decades. In future research it will thus be crucial to establish whether other insect species also show a high rate of recent TE invasions (e.g. using museomics [Raxworthy and Smith, 2021]) and to continue to monitor natural populations for novel invasions with community efforts such as DrosEU [Kapun et al., 2021]. Furthermore, it will be important to identify risk factors contributing to elevated rates of TE invasions (e.g. ecological factors or idiosyncrasies in the host defence). Finally, it will be crucial to estimate the phenotypic effects of these TE invasions, especially given the substantial impact of TEs on the evolution of genomes and phenotypes.

## Materials and methods

### Detecting the three new transposon invasions

To identify candidates of TEs that may have invaded *D. melanogaster* recently, we used our new tool GenomeDelta (Pianezza et al, in preparation). GenomeDelta is based on the idea that recent TE invasion will lead to sequences present in young strains (i.e. collected after the invasion) that are absent in old strains (collected before the invasion). GenomeDelta therefore aims to identify sequences that are present in young but absent in old strains. Briefly, GenomeDelta, aligns short-reads of an old strain to the genome assembly of a young strain, identifies gaps in the coverage as for example resulting from insertions of a novel TE, extracts the sequences responsible for the coverage gaps and clusters them based on homology. We discovered the invasions of *MLE, Souslik* and *Transib1* by aligning short-read data of three old strains (SRR23876563, SRR11846565, SRR11846560; collected between 1810 - 1965) to the assemblies of 4 recently collected strains (GCA 020141595.1, GCA 020141505.1, GCA 020141485.1, GCA 020141515.1; collected between 2015 - 2016).

### Characterising the invasions

We generated a phylogenetic tree based on the reverse transcriptase domain of the 7 LTR retrotransposons as described previously [Pianezza et al., 2023]. Briefly, we picked several sequences from each of the known LTR superfamily/groups [Kapitonov and Jurka, 2003, Quesneville et al., 2005], performed a blastx search to identify the reverse transcriptase domain [Wheeler et al., 2007], created a multiple sequence alignment with MUSCLE (v3.8.1551)Edgar [2004], generated trees with BEAST (v2.7.5)Bouckaert et al. [2019] and extracted the maximum credibility tree with TreeAnnotator (v2.7.5)Bouckaert et al. [2019].

We investigated the population frequency of the 11 TEs and the diverged fragments using short-reads from a pooled population collected in Ukraine in 2016 (SRR8494428) [Kapun et al., 2021]. Diverged fragments were identified with RepeatMasker [Smit et al., 2013-2015] using a minimum length threshold of 300bp and a maximum divergence of 20. The insertions frequencies were estimated using PopoolationTE2 [Kofler et al., 2016] (–min-distance -200 –max-distance 300 –min-count 2 –max-otherte-count 2 –max-structvar-count 2).

### Copy number estimates using short read data

We investigated the abundance of the 11 TEs in multiple publicly available short-read data sets. In total we analysed 585 strains of *D. melanogaster* [Grenier et al., 2015, Schwarz et al., 2021, Long et al., 2013, Lange et al., 2021, Rech et al., 2022, Kapun et al., 2021, Shpak et al., 2023, Pool et al., 2012]. For an overview of all analysed data see supplementary file S1. To estimate the copy numbers of the TE in the samples we used our tool DeviaTE [Weilguny and Kofler, 2019]. To do this, we aligned the reads to a database consisting of the consensus sequences of the 11 TEs that invaded during the last 200 years, and three single copy genes (*Antennapedia, Osi6* and *Wingless*; with bwa bwasw (version 0.7.17-r1188) [Li and Durbin, 2009]. DeviaTE then estimates the copy number of a TE (e.g. *Souslik*) by normalizing the coverage of the TE by the coverage of the single copy genes. We also used DeviaTE to visualize the abundance and diversity of the TEs as well as to compute the frequency of SNPs in TEs.

### Copy number estimates in long-read assemblies

We investigated the abundance of the 11 TEs in long-read assemblies. The 49 analysed *D. melanogaster* strains were described previously [Pianezza et al., 2023] (data from [Chakraborty et al., 2019, Wierzbicki et al., 2021, Hoskins et al., 2015, Rech et al., 2022]). For an overview of all analysed assemblies see supplementary file S2. We identified insertions in these assemblies using RepeatMasker (open-4.0.7; -no-is -s -nolow)[Smit et al., 2013-2015].

### Identification of diagnostic SNPs

To identify SNPs that are diagnostic for canonical insertions, we investigated the frequencies of alleles in the TE sequences. For example, if 10 insertions of a TE have the allele ‘A’ at position 243 of the TE sequence and 5 the allele ‘T’ frequencies of the alleles at the given site are are 10*/*15 and 5*/*15 for ‘A’ and ‘T’, respectively. The output of DeviaTE [Weilguny and Kofler, 2019] was used to obtain the frequency of SNPs. We aimed to identify alleles of the TE sequence that are present in canonical but absent in degraded insertions (i.e. diagnostic for canonicial insertions), by comparing the allele frequencies between an old (SRR8494428; collected in 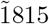) and a young sample (SRR23876563; collected in 2016). We considered a SNP to be diagnostic for canonical insertions if the minor allele frequency was *>* 0.1 in the young and *>* 0.95 in the old sample. With this approach, we identified multiple diagnostic SNPs for most of the 11 TEs that invaded *D. melanogaster* during the last century. Finally, we averaged the frequencies of all diagnostic SNPs for a given TE family. No reads aligned to the *P* -element and *Opus* in the old samples, preventing the calculation of allele frequencies in these samples. For these two TEs we used frequencies of 1.0 or 0.0 to indicate the presence and absence of reads aligning to the TE, respectively. We used the diagnostic SNPs (and the copy number of the TE) to trace the invasions in the short read data.

### Origin of horizontal transfer

To identify the origin of the horizontal transfer that triggered the 11 invasions we obtained long-read assemblies of 266 drosophilids [Kim et al., 2021, 2023] and of 1225 arthropod reference genomes from NCBI. The arthropod genomes were found by filtering for “arthropoda”, “chromosome level” and “reference” at the NCBI database. The list of all analysed drosophilid and arthropod species, including the source, can be found in supplementary files S4, S3. We used RepeatMasker [Smit et al., 2013-2015] (open-4.0.7; -no-is -s -nolow) to identify sequences with similarity to the 11 TEs that invaded *D. melanogaster* recently in these genomes. Using a Python script, we identified the best hit for each TE in each assembly (i.e. the highest alignment score) and than estimated the similarity between this best hit and the TE using the equation *s* = *rms*_*best*_*/rms*_*max*_, where *rms*_*best*_ is the highest RepeatMasker score (rms) in a given assembly and *rms*_*max*_ the highest score in any of the analysed assemblies. A *s* = 0 indicates no similarity to the consensus sequence of the TE, whereas *s* = 1 represent the highest possible similarity.

To generate a phylogenetic trees for the TEs, we extracted the sequences of full-length insertions (*>* 80% of the length) from species having at least one full-length insertion using bedtools [Quinlan and Hall, 2010](v2.30.0). For each TE, a multiple sequence alignment of the insertions was generated with MUSCLE (v3.8.1551) [Edgar, 2004] and a tree was generated with BEAST (v2.7.5) [Bouckaert et al., 2019].

## Acknowledgments

We thank thank all members of the Institute of Population Genetics for feedback and support. SS would like to thank F. Belt, S. & F. Emery, and P. Senn for inspiration during the production of this manuscript.

## Author contributions

RP discovered the three new TE invasions. RP, AS, SS, RK conceived the work. RP, AS and AH analysed the data. SS and RK wrote the manuscript. RP and AS contributed to writing. All authors read and approved the manuscript

## Data Availability

All analyses performed in this work were documented in RMarkdown and have been made publicly available, together with the resulting figures, at GitHub https://github.com/rpianezza/Dmel-200years.

