## Supplementary info for "Unveiling the complete invasion history of *D. melanogaster*: three horizontal transfers of transposable elements in the last 30 years"

### Supplementary figures and tables

#### Supplementary figures

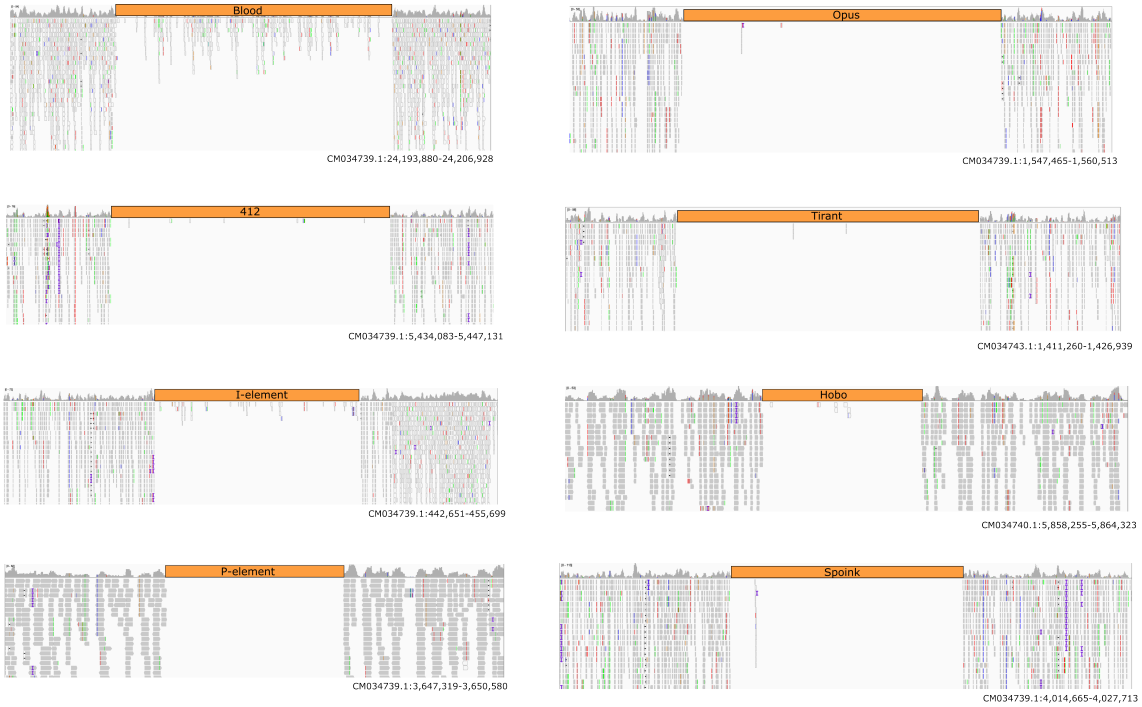


Figure 1: Example of coverage gaps caused by the 8 TE families previously shown to have invaded *D. melanogaster* populations within the last two centuries: *Blood*, *Opus*, *412*, *Tirant*, the *I*-element, *Hobo*, the *P*-element and *Spoink*. Short reads of H10 (∼1810; [Shpak et al., 2023]) were aligned to the assembly of TOM008 (2016; [Rech et al., 2022]).

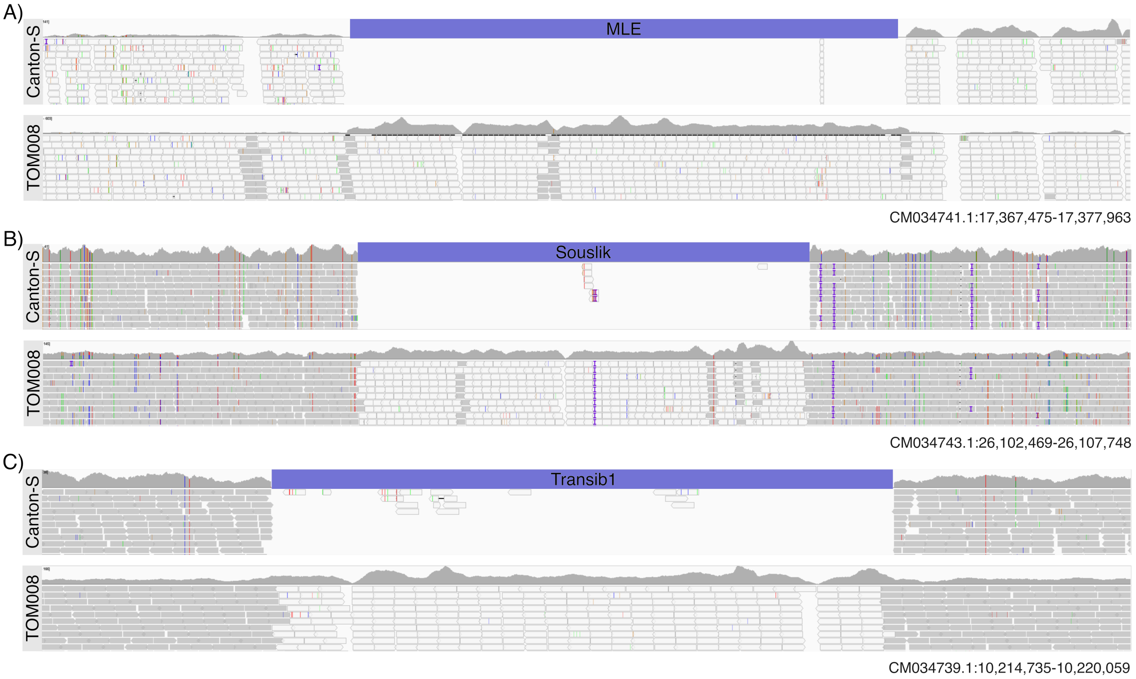


Figure 2: Example of coverage gaps caused by *MLE* (A) *Souslik* (B) and *Transib1* (C) insertions. Short reads of Canton-S (1935; [Wierzbicki et al., 2021]) were aligned to the assembly of TOM008 (2016; [Rech et al., 2022]). As control we also aligned the short-reads of TOM008 to the assembly of the same strain.

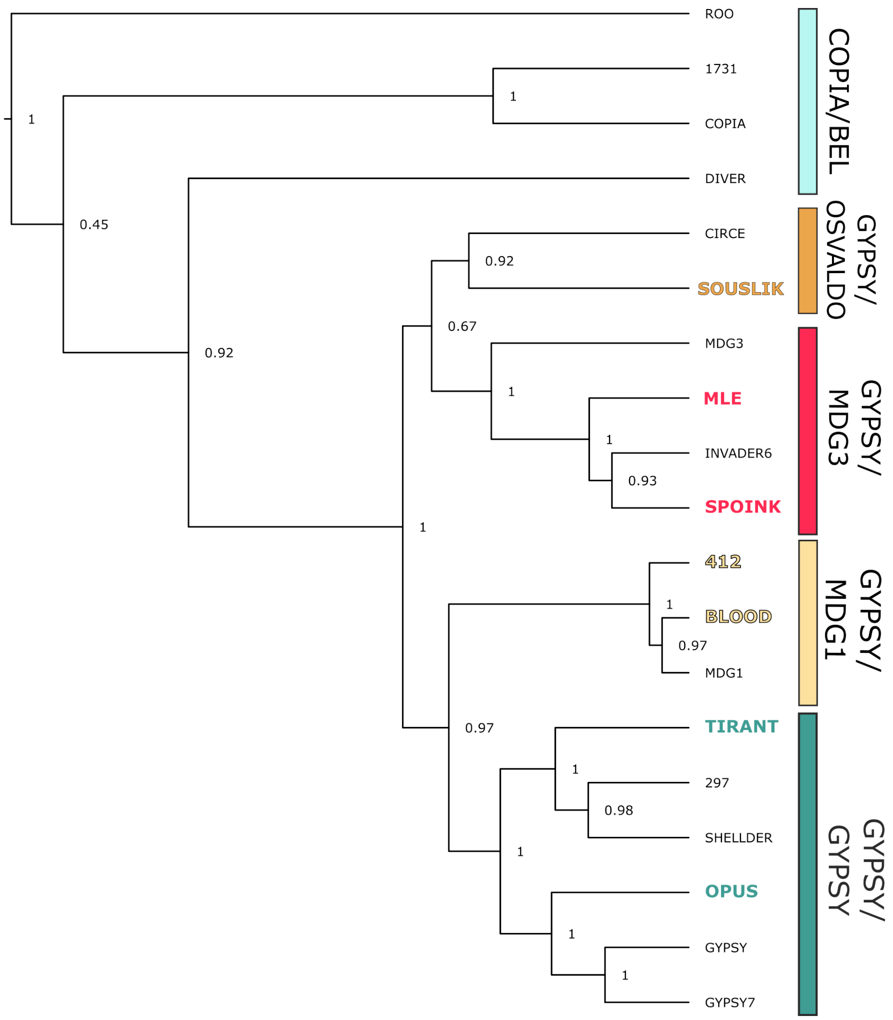


Figure 3: Phylogenetic tree of *Souslik* and *MLE* based on the reverse-transcriptase domain of *pol*. As reference we picked some families for each of the main superfamilies/groups of LTR retrotransposons and included other TEs that recently invaded *D. melanogaster* (*Tirant*, *412*, *Blood*, *Opus*) or *D. simulans* (*Shellder*) [Kapitonov and Jurka, 2003, Scarpa et al., 2023, Pianezza et al., 2023, Ding et al., 2016]. Our data suggest that *MLE* is a member of the *gypsy/mdg3* group and *Souslik* is a member of the *gypsy/osvaldo* group.

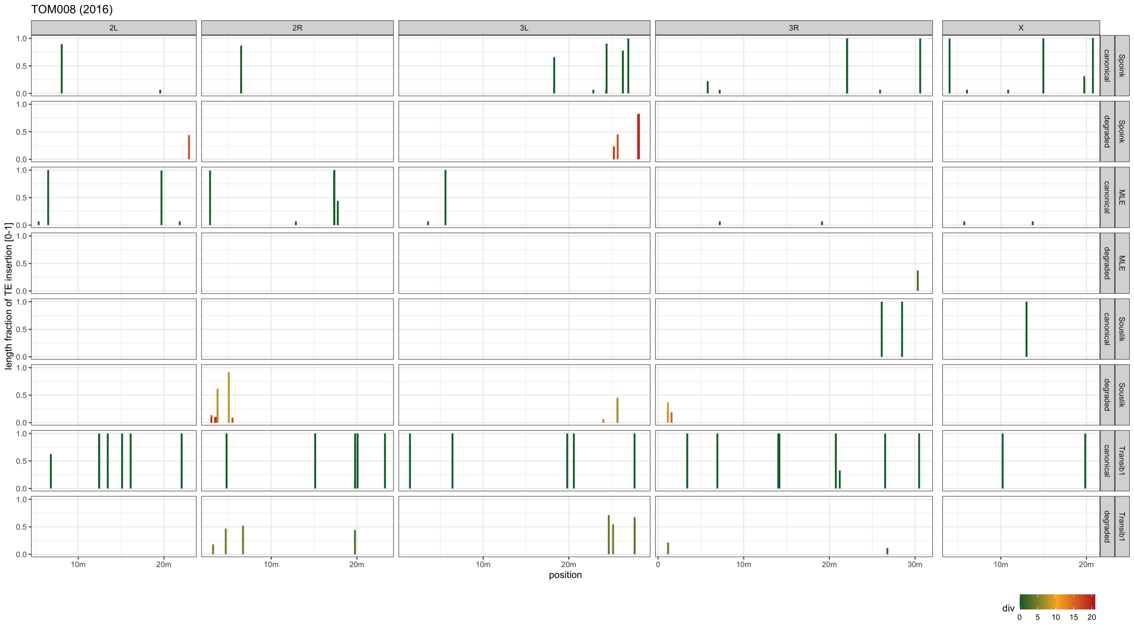


Figure 4: Overview of *Spoink*, *MLE*, *Souslik* and *Transib1* insertions in a long-read assembly of *TOM008* (identified with RepeatMasker; divergence $< 20\%$). Plots show the genomic position and the length of fragments matching with the consensus sequence of a given TE. Colors represent divergence from the consensus sequence. Based on a threshold of 1.5%, we classified the fragments into canonical ($\leq 1.5\%$) and degraded ($> 1.5\%$). The length of the TE (y-axis) is normalized to the full-length element. Note that the diverged insertions of the TEs are close to the ends of the chromosome arms (likely heterochromatic regions).

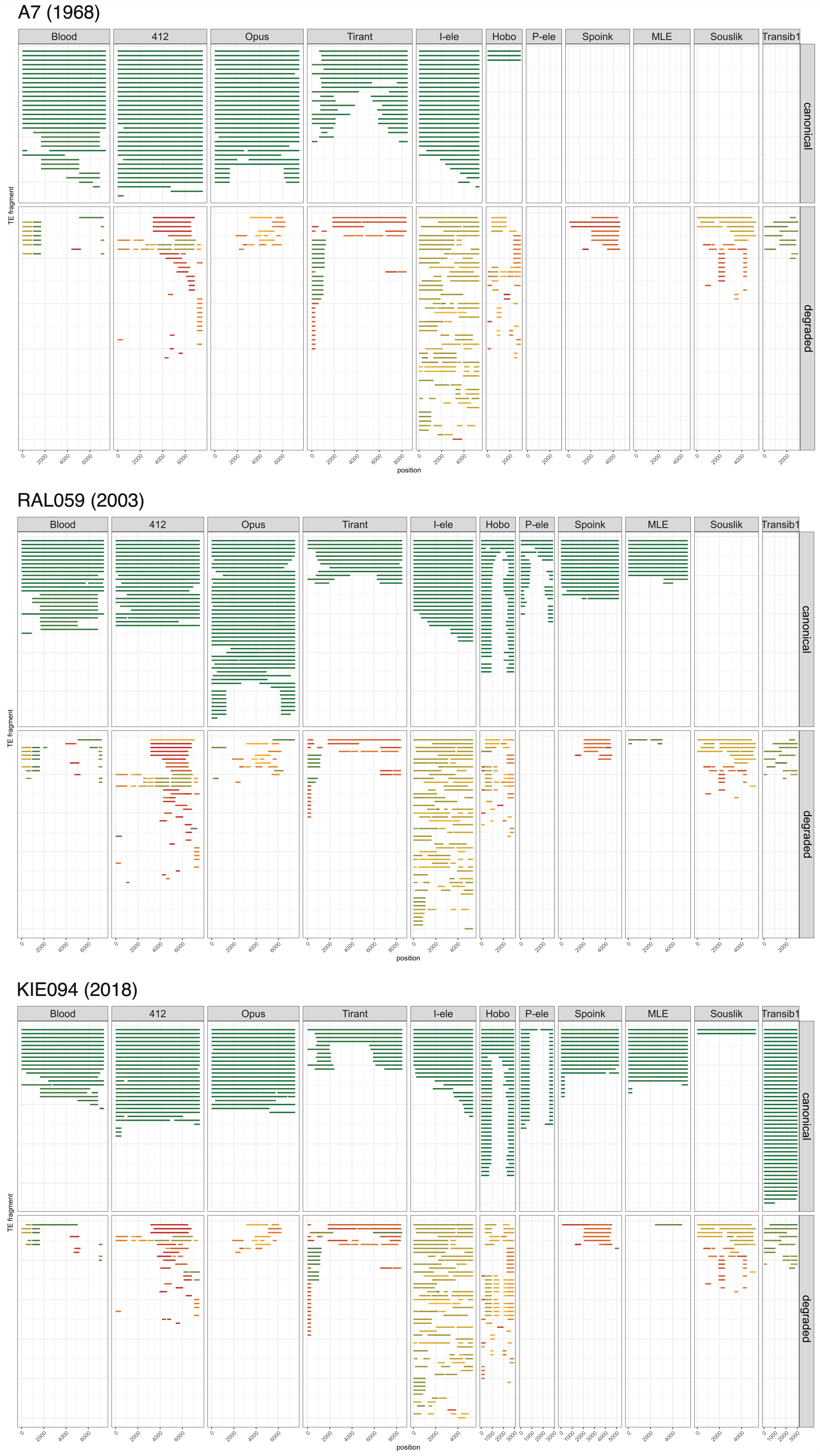


Figure 5: Regions matching with the consensus sequence of a TE in assemblies of strains collected between 1968 and 2018. Results are shown separately for canonical (≤1.5% divergence) and degraded (>1.5% divergence) insertions. The number of TE families with canonical insertions increases with the sampling date of strains (the *P*-element, *Spoink*, *MLE*, *Souslik* and *Transib1*).

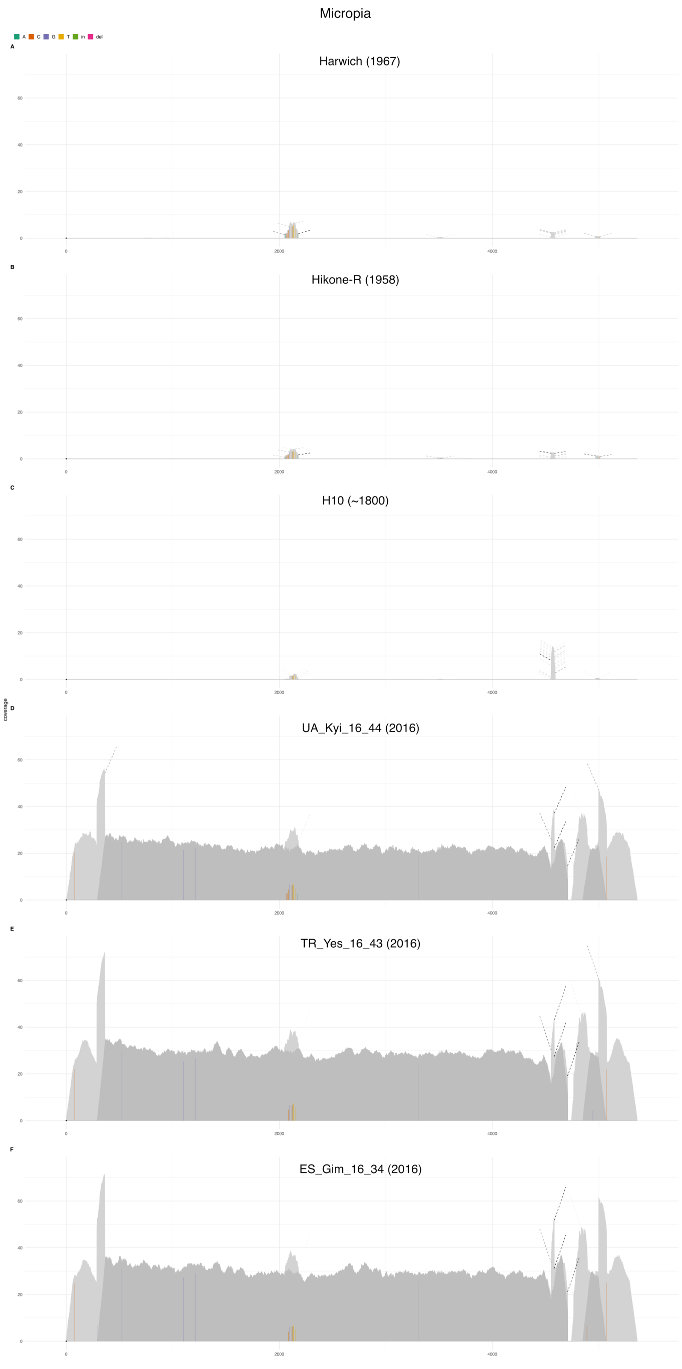


Figure 6: DeviaTE plots of *MLE* for six *D. melanogaster* strains collected during the last centuries. The short reads were aligned to the consensus sequence of TEs and the coverage was normalized to the coverage of single-copy genes. SNPs and indels are shown as colored lines. Coverage based on unambiguously and ambiguously aligned reads is shown in dark and light grey, respectively. Note that very few reads of old strains ($\leq$ 1967) align to *MLE* whereas a contiguous coverage is observed for more recently collected strains (2016).

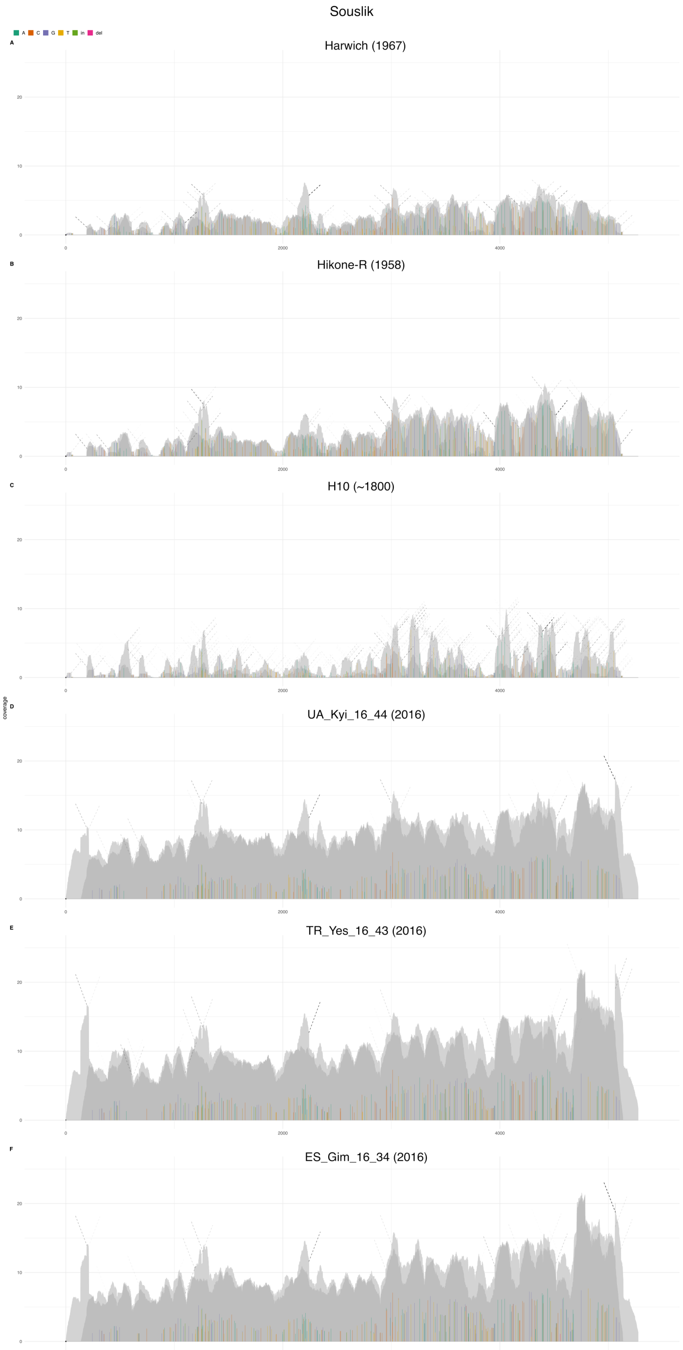


Figure 7: DeviaTE plots of *Souslik* for six *D. melanogaster* strains collected during the last centuries. The short reads were aligned to the consensus sequence of TEs and the coverage was normalized to the coverage of single-copy genes. SNPs and indels are shown as colored lines. Coverage based on unambiguously and ambiguously aligned reads is shown in dark and light grey, respectively. Note that in old strains ($\leq$ 1967) solely a few diverged reads align to *Souslik*, whereas a high contiguous coverage is observed for more recently collected strains (2016). Furthermore, the increase in coverage is largely due to reads having a high similarity to the consensus sequence of *Souslik*.

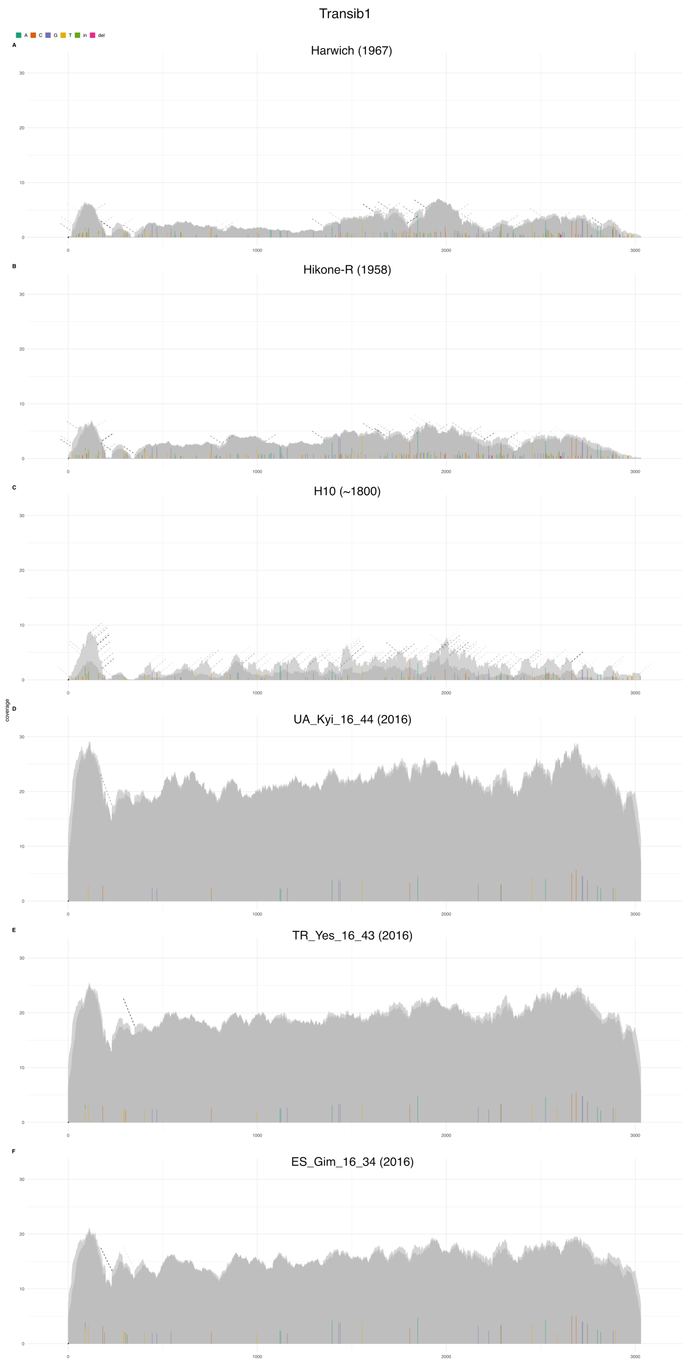


Figure 8: DeviaTE plots of *Transib1* for six *D. melanogaster* strains collected during the last centuries. The short reads were aligned to the consensus sequence of TEs and the coverage was normalized to the coverage of single-copy genes. SNPs and indels are shown as colored lines. Coverage based on unambiguously and ambiguously aligned reads is shown in dark and light grey, respectively. Note that in old strains ($\leq$ 1967) solely a few diverged reads align to *Transib1*, whereas a high contiguous coverage is observed for more recently collected strains (2016). Furthermore the increase in coverage is largely due to reads having a high similarity to the consensus sequence of *Transib1*.

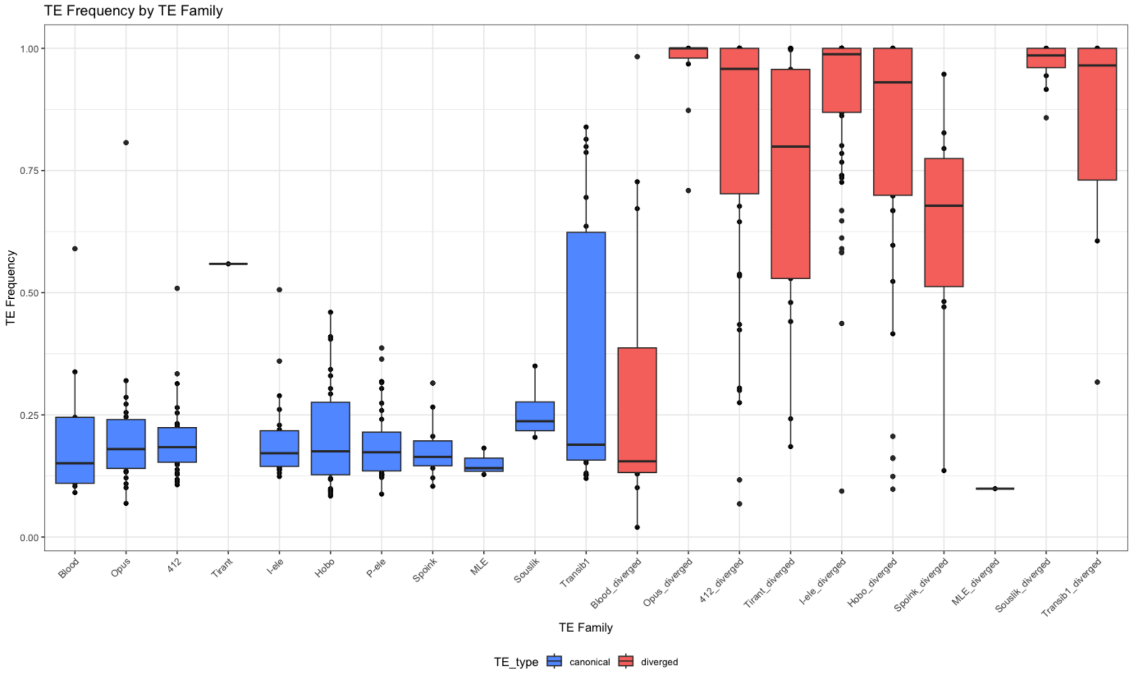


Figure 9: Population frequency of canonical (blue) and diverged (red) insertions for the 11 TEs that invaded *D. melanogaster* during the last two centuries. Population frequencies were estimated with PoPoolationTE2 based on pooled short-read data from a population collected 2016 in Ukraine (SRR8494428) [Kapun et al., 2021, Kofler et al., 2016]. Canonical insertions are largely segregating at a low frequency, consistent with a recent invasion of the canonical TEs. By contrast, diverged insertions are mostly fixed, suggesting a more ancient origin. Three exceptions may be worth mentioning. For the *P*-element no diverged insertions can be found. For *MLE* we solely found a single diverged insertion, which however likely arose during the recent *MLE* invasion (see main manuscript fig 1B; hence it is likely a very recent insertion). Based on the divergence between the two LTRs, the diverged *Blood* insertions have an estimated age of about 650.000 years [Scarpa et al., 2023] and should thus largely be fixed in the population. Possible explanations for the observed low population frequency of diverged *Blood* insertions may be biological (e.g. recent structural rearrangements, delayed fixation) or technical (e.g. problems with estimating the population frequency of highly similar insertions; canonical and diverged *Blood* have a similar sequence [Scarpa et al., 2023]).

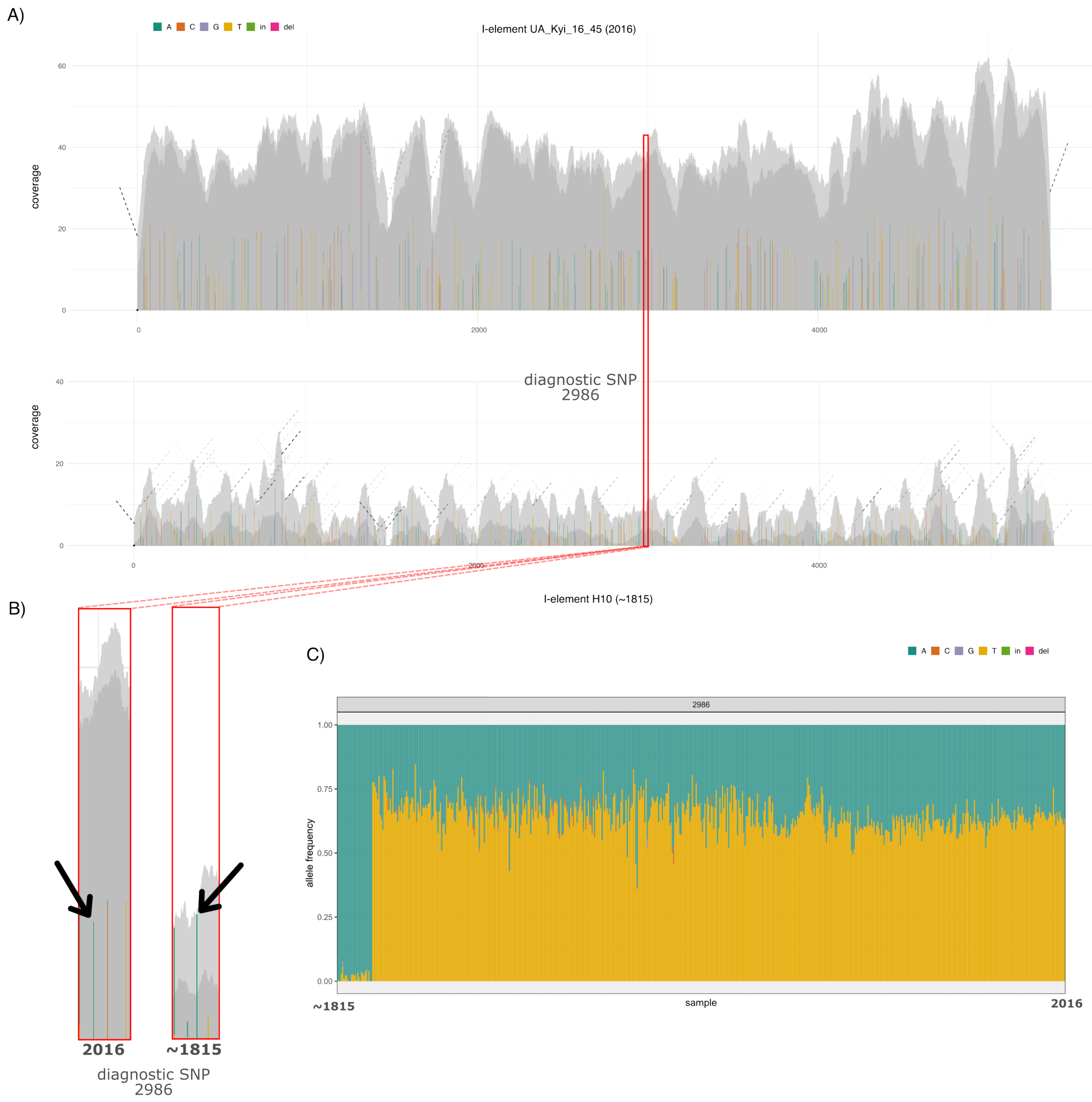


Figure 10: Diagnostic SNPs enable tracing the invasion of canonical insertions. A) DeviaTE plots for a strain collected before (∼1815) and after (2016) the invasion of the canonical *I*-element. Colored lines indicate the abundance of SNPs and indels (the reference allele is not shown). Many SNPs are segregating in the sample from 2016 (the presence of canonical and degraded insertions leads to two alleles) but are fixed in the sample from 1815 (the degraded insertions have just one allele). B) Magnification of one of these SNPs, at position 2986 of the I-element. Arrows indicate the same site. This SNP has two alleles in the sample collected in 2016, but just one in the sample collected in ∼1815. Hence, the reference allele of this SNP is diagnostic for the canonical *I*-element. C) Frequencies of the two alleles at the diagnostic SNP at position 2986 of the *I*-element in 585 *D. melanogaster* strains collected during the last two centuries. A sudden shift in the allele frequencies is observed at the time of the invasion of the canonical *I*-element. The frequency of the SNP at position 2986 can thus be used to trace the invasion of the canonical *I*-element.

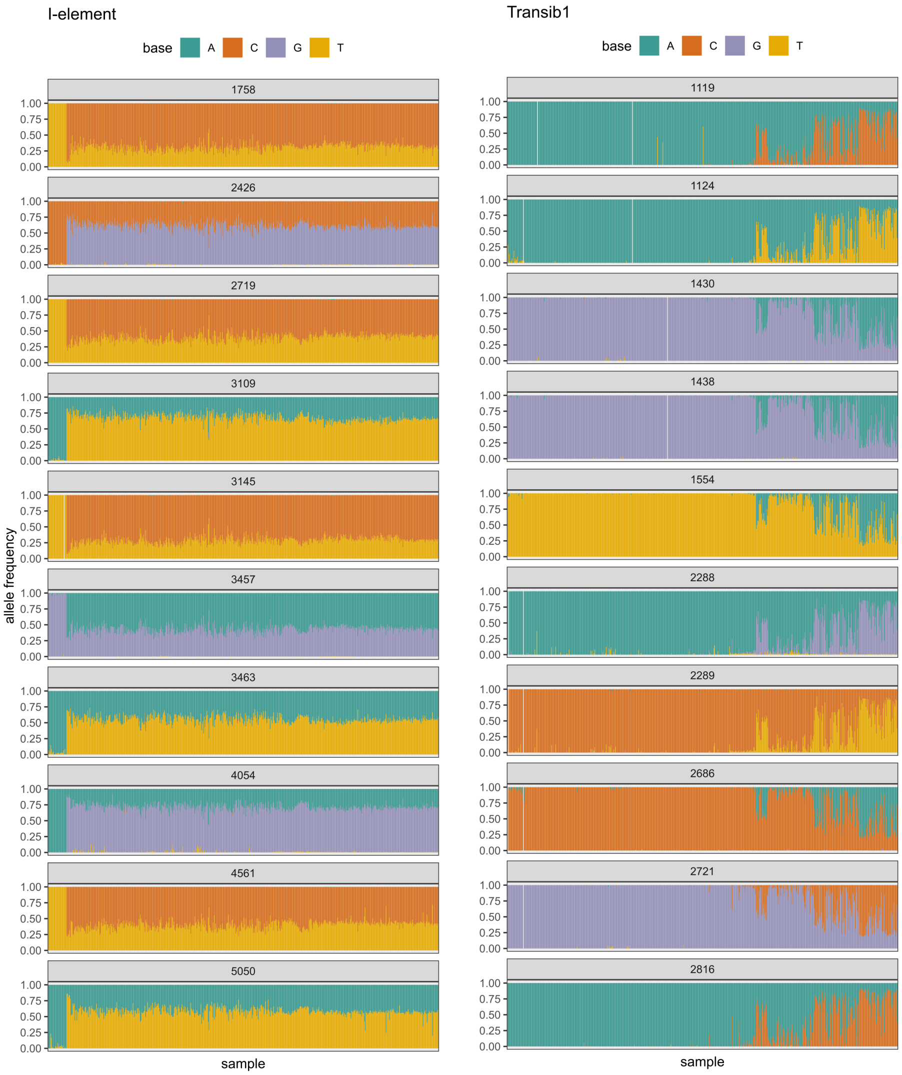


Figure 11: Diagnostic SNPs enable to consistently infer the timing of the invasion. We show the allele frequency of several SNPs that are diagnostic for canonical insertions of the *I*-element and *Transib1*. Allele frequencies are shown for 585 strains sampled during the last two centuries (ascending order by year). For each TE, ten randomly chosen diagnostic SNPs are shown (the position of the SNP in the consensus sequence is shown in the top panel). A consistent shift in the allele frequency can be observed for all diagnostic SNPs at the onset of the invasion of canonical insertions.

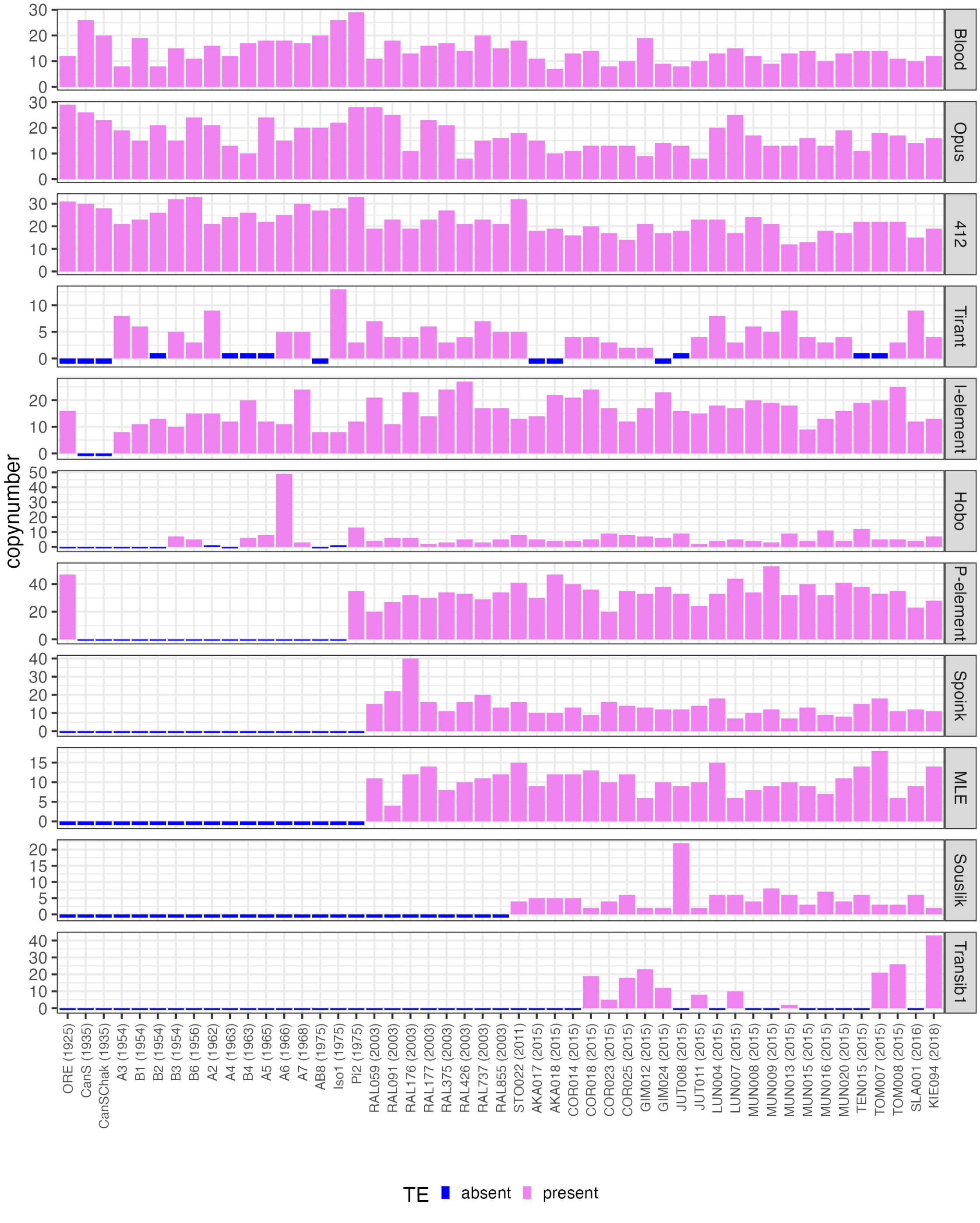


Figure 12: Abundance of canonical insertions for the 11 TEs that invaded *D. melanogaster* recently in long-read assemblies of different *D. melanogaster* strains collected between 1925 and 2018. We assumed that canonical insertions have <1.5% divergence from the consensus sequence over > 80% of the length, except for the *P*-element and *Hobo*, where we used a length threshold of 50%, to account for insertions with internal deletions [Black et al., 1987].

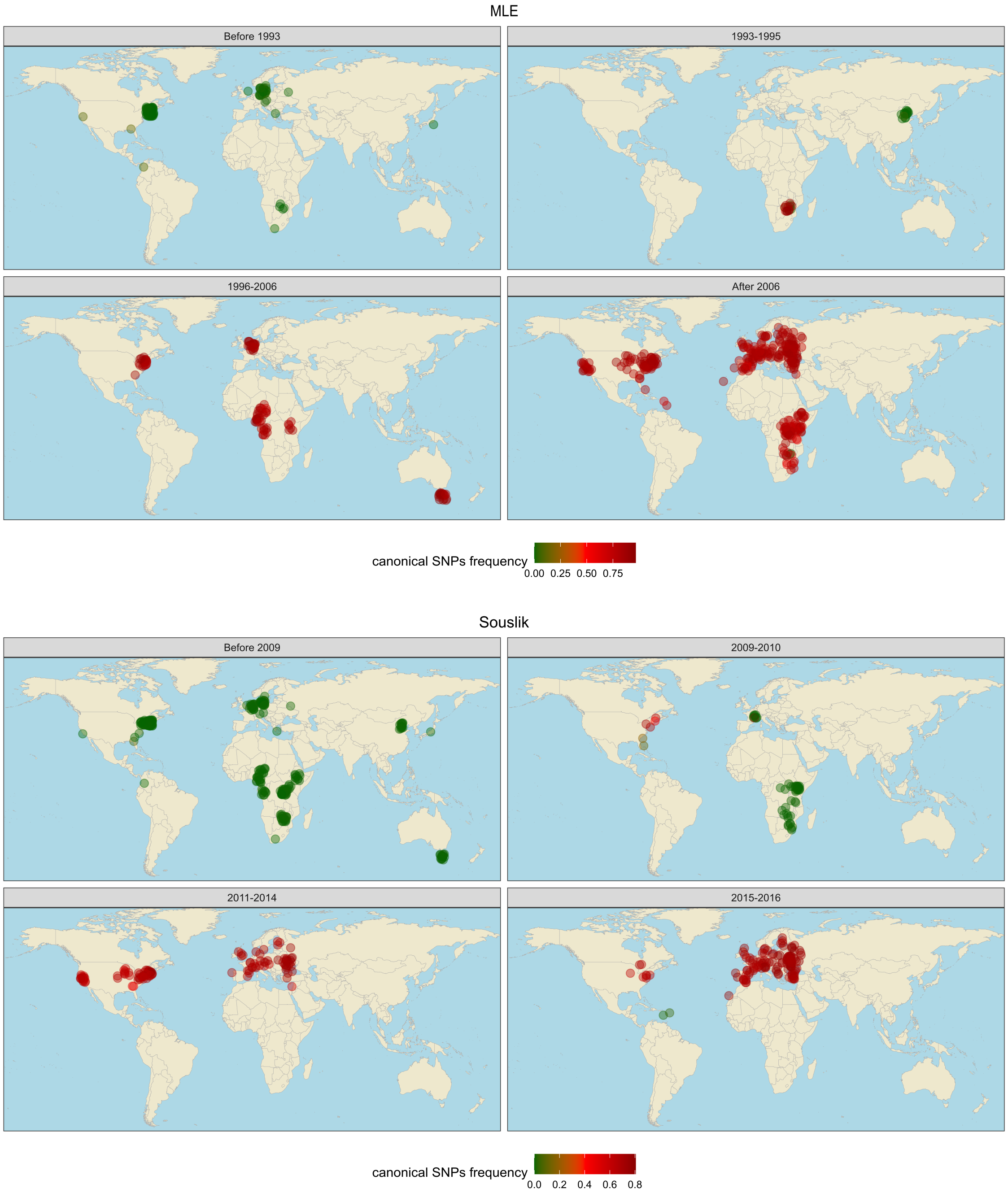


Figure 13: Geographic spread of canonical *MLE* and *Souslik* insertions in *D. melanogaster* samples (strains and pooled populations) collected at different geographic locations during the last decades. Colors of dots refers to the frequency of SNPs diagnostic for canonical insertions.

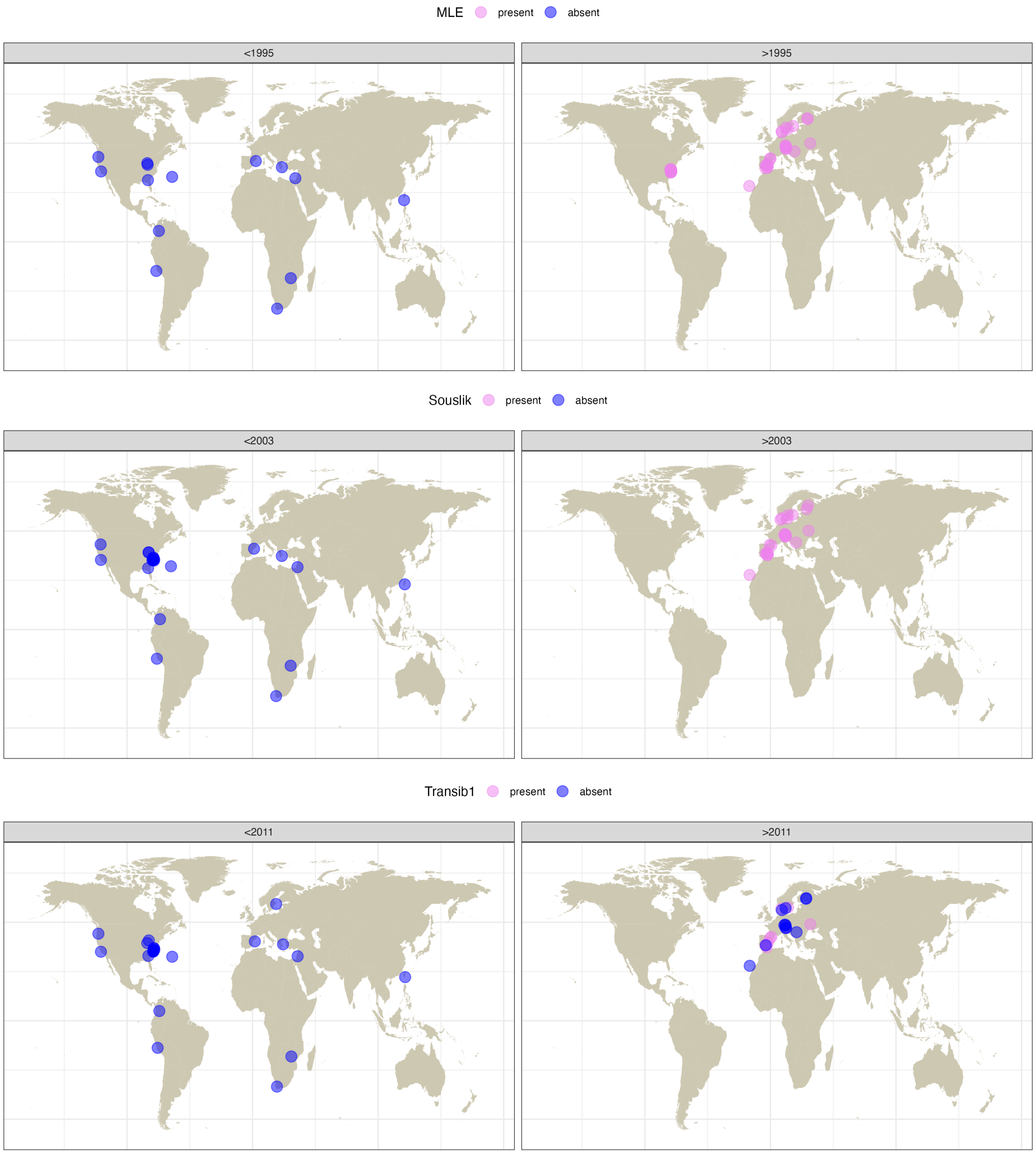


Figure 14: Geographic spread of canonical *MLE*, *Souslik* and *Transib1* insertions in long-read assemblies of *D. melanogaster* strains collected at different geographic locations during the last decades. The number of canonical insertions in the assemblies was estimated with RepeatMasker (<1.5% divergence; >80% of length). absent: number of canonical insertions is zero; present: at least one canonical insertion was found

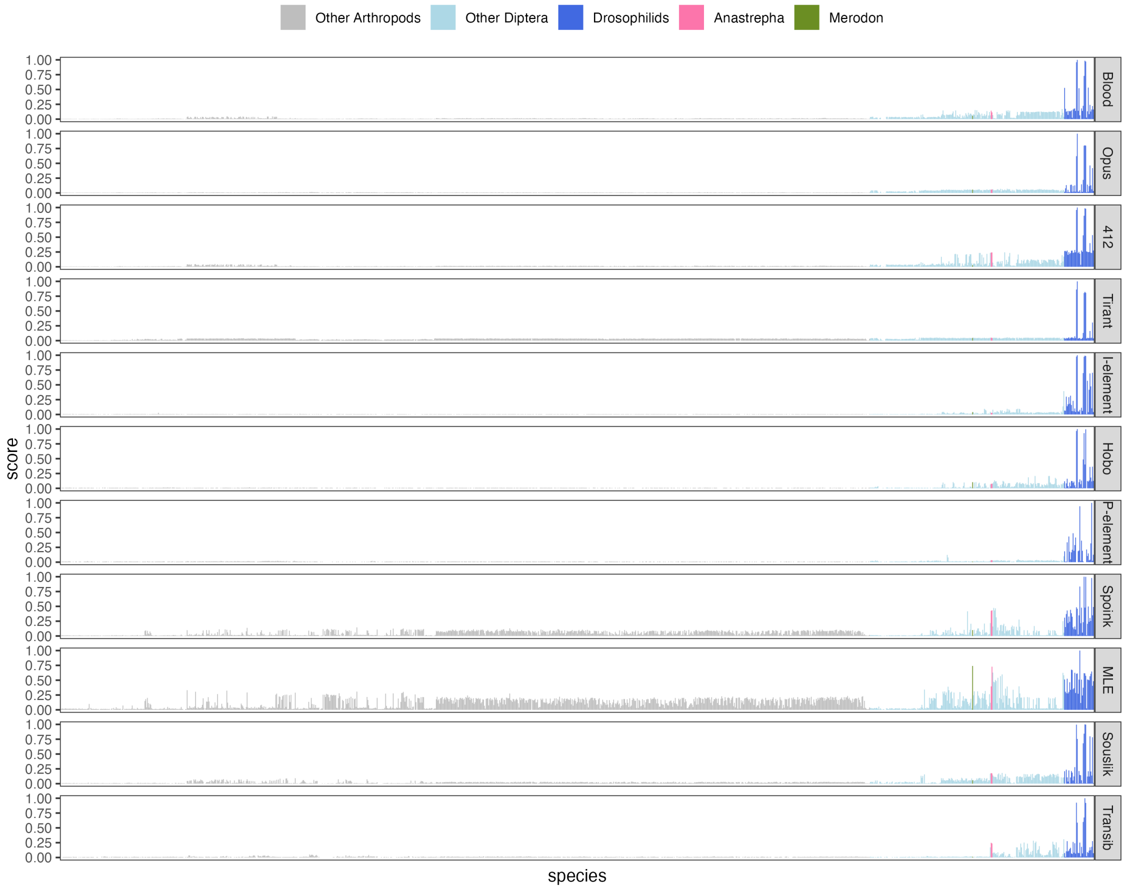


Figure 15: Similarity of the consensus sequence of the 11 TEs which spread in *D. melanogaster* recently, with TE insertions in 1226 assemblies of arthropod species. The barplots show, for each species, the similarity between the given TE and the best match in an assembly. For example, a value of 0.9 indicates that at least one insertion in an assembly has a high similarity to the given TE. For the 11 TE families, few similar sequences can be found outside of the *drosophilids*, with the exception of *MLE*, where we found similar sequences in *Anastrepha obliqua* and *Merodon equestris*

### Supplementary tables

Table 1: TE content in long-read assemblies of recently collected *D. melanogaster* strains. For each of the 11 TEs that invaded *D. melanogaster* populations recently, we show the genomic proportion in base pairs. The assembly size (genome length) and the genomic fraction (in percent) occupied by these TEs are also shown.

| TE | TOM008 | SLA001 | KIE094 | COR025 |
|---|---|---|---|---|
| *412* | 206,031 | 154,672 | 193,658 | 128,098 |
| *Blood* | 101,936 | 140,926 | 132,474 | 124,489 |
| *Hobo* | 71,924 | 60,462 | 70,052 | 69,615 |
| *I*-element | 160,639 | 83,447 | 84,785 | 98,348 |
| *MLE* | 38,514 | 52,407 | 79,215 | 78,998 |
| *Opus* | 181,020 | 186,349 | 166,319 | 170,726 |
| *P*-element | 41,862 | 27,891 | 32,545 | 47,439 |
| *Souslik* | 15,862 | 41,436 | 10,563 | 35,415 |
| *Spoink* | 75,024 | 71,468 | 65,000 | 83,461 |
| *Tirant* | 40,188 | 97,229 | 60,994 | 90,723 |
| *Transib1* | 78,997 | 11,408 | 134,160 | 55,035 |
| total | 1,011,997 | 927,695 | 1,029,765 | 982,347 |
| genome length | 110,897,714 | 116,372,588 | 116,565,937 | 116,374,970 |
| new TE fraction | 0.91% | 0.80% | 0.88% | 0.84% |
